# Deep profiling of antigen-specific B cells from different pathogens identifies novel compartments in the IgG memory B cell and antibody-secreting cell lineages

**DOI:** 10.1101/2023.12.19.572339

**Authors:** M. Claireaux, G. Elias, G. Kerster, LH. Kuijper, MC. Duurland, AGA. Paul, JA. Burger, M. Poniman, W. Olijhoek, N. de Jong, R. de Jongh, E. Wynberg, HDG. van Willigen, M. Prins, GJ. De Bree, MD. de Jong, TW. Kuijpers, F. Eftimov, CE. van der Schoot, T. Rispens, JJ. Garcia-Vallejo, A. ten Brinke, MJ. van Gils, SM. van Ham

## Abstract

A better understanding of the bifurcation of human B cell differentiation into memory B cells (MBC) and antibody-secreting cells (ASC) and identification of MBC and ASC precursors is crucial to optimize vaccination strategies or block undesired antibody responses. To unravel the dynamics of antigen-induced B cell responses, we compared circulating B cells reactive to SARS-CoV-2 (Spike, RBD and Nucleocapsid) in COVID-19 convalescent individuals to B cells specific to Influenza-HA, RSV-F and TT, induced much longer ago. High-dimensional spectral flow cytometry indicated that the decision point between ASC- and MBC-formation lies in the CD43+CD71+IgG+ Activated B cell compartment, showing properties indicative of recent germinal center activity and recent antigen encounter. Within this Activated B cells compartment, CD86+ B cells exhibited close phenotypical similarity with ASC, while CD86− B cells were closely related to IgG+ MBCs. Additionally, different activation stages of the IgG+ MBC compartment could be further elucidated. The expression of CD73 and CD24, regulators of survival and cellular metabolic quiescence, discerned activated MBCs from resting MBCs. Activated MBCs (CD73-CD24lo) exhibited phenotypical similarities with CD86− IgG+ Activated B cells and were restricted to SARS-CoV-2 specificities, contrasting with the resting MBC compartment (CD73-/CD24hi) that exclusively encompassed antigen-specific B cells established long ago. Overall, these findings identify novel stages for IgG+ MBC and ASC formation and bring us closer in defining the decision point for MBC or ASC differentiation.

**Importance:** In this study, researchers aimed to better understand human B cell differentiation and their role in establishing long-lived humoral immunity. Using high-dimensional flow cytometry, they studied B cells reactive to three SARS-CoV-2 antigens in individuals convalescent for COVID-19, and compared their phenotypes to B cells reactive to three distinct protein antigens derived from vaccines or viruses encountered months to decades before. Their findings showed that Activated B cells reflect recent germinal center graduates that may have diverse fates; with some feeding the pool of antibody-secreting cells and others fueling the resting memory B cell compartment. Activated B cells gradually differentiate into resting memory B cells through an activated MBC phase. Increased expression of the cellular metabolic regulators CD73 and CD24 in resting memory B cells distinguishes them from the activated memory B cells phase, and is likely involved in sustaining a durable memory of humoral immunity. These findings are crucial for the development of vaccines that provide lifelong protection and may show potential to define reactive B cells in diseases where the cognate-antigen is still unknown such as in autoimmunity, cancers, or novel viral outbreaks.

## INTRODUCTION

Formation of long-lived antibody-secreting cells (ASC) and memory B cells (MBC) is essential to generate and maintain protective humoral immunity against invading pathogens. After infection or vaccination, in secondary lymphoid organs, naive B cells that recognize their cognate antigens can participate in either extrafollicular (EF) or germinal center (GC) responses. The EF pathway (*1*) induces early MBCs and short-lived ASCs that mostly harbor an IgM isotype with limited B cell receptor (BCR) mutations (*2*). Short-lived ASCs contribute to the rapid production of antibodies of modest affinity, while early MBCs can participate in a secondary response (*3*). In contrast, with help of cognate follicular T helper cells (Tfh), B cells that follow the GC pathway undergo affinity maturation and class switching to generate high affinity IgG, IgA, and IgE B cells. Briefly, responsive GC B cells migrate to GC Dark zones where they undergo rapid proliferation and experience somatic hypermutations, resulting in B cells with slightly mutated BCRs. Subsequent migration from the Dark Zones to Light Zones of the GC enables the newly generated B cells to re-encounter antigens presented by follicular dendritic cells. B cells expressing BCRs with greater affinity than their counterpart will successfully capture more antigens and re-acquire cognate Tfh help for further rounds of proliferation and mutation. This process ultimately leads to development of class-switched high-affinity GC B cell clones that can further differentiate in MBCs, and long-lived plasma cells (LLPCs) that migrate away and encapsulate into the bone marrow (*4*). Understanding the underlying mechanisms that dictate the differentiation of B cells into either MBCs or LLPCs is a key focus in the B cell research field. LLPCs are morphologically, transcriptionally, and metabolically configured to secrete antibodies from the bone marrow into the circulation and provide long-lasting humoral immunity (*3*), while MBCs can either rapidly proliferate and differentiate into short-lived ASC or migrate to the GC for a new round of somatic hypermutations and affinity selection (*5*). As a result, following a viral infection or vaccination, an efficient GC reaction provides a durable immune response with improved neutralization breadth and potency. This limits the range of escape options for the pathogen and prepares the host for future encounters with pathogen variants. Vaccine strategies are designed to preferentially promote GC over EF pathway. In spite of that, there is still a lack of knowledge in the phenotype and ontology of circulating B cells that could serve as biomarkers of an optimal protective and durable immunity, namely experienced B cells that egress from ongoing GC reactions within lymph nodes, MBCs that display long-lived properties, and MBCs that would preferentially participate in a secondary GC response. In this study, we aim to delineate heterogeneities in circulating antigen-specific B cell compartments and relate differences to time of exposure to the antigen (recent versus longer ago) to identify quiescent MBCs and recently antigen-activated B cells.

Circulating B cells have traditionally been separated based on expression of IgD and CD27 surface proteins into four subsets: IgD+ CD27− (naïve), IgD+ CD27+ (unswitched MBCs), IgD− CD27− (double negative (DN) MBCs), and IgD− CD27+ (classical MBCs) (*6*). Initially, CD27+ B cells were considered the sole MBC compartment, but later DN B cells were shown to also display memory features. DN B cells can be recalled following stimulation and, while less mutated than CD27+ B cells, there is still some clonal overlap between the two compartments (*7*). Recently, there has been renewed interest in the CD45RB marker, which has been proposed as a broader MBC marker to distinguish early canonical B cells from DN B cells within the IgD− CD27− population (*8*, *9*). In addition, there is a consensus in the use of CD21 marker to further classify circulating B cells: IgD+ CD21+ CD27− (Naïve), IgD− CD21+ CD27+ (classical MBCs), IgD− CD21− CD27− (DN2/DN3), IgD− CD21+ CD27− (DN1) and IgD− CD21− CD27+ (Activated B cells, ActBCs). DN2 and DN3 can be further separated based on expression of CD11c and T-Bet (DN2, CD11c+ T-Bet+; DN3 CD11c− T-Bet−) (*10*, *11*). In chronic infection, DN B cells were shown to accumulate and display exhausted properties associated with an inhibitory phenotype (*12*, *13*). However, in systemic lupus erythematosus (SLE) (*14*) and COVID-19 (*11*, *15*), DN2 and DN3 B cells respectively are active and represent the early EF B cell response that may be poised for ASC differentiation. In contrast, CD27+ ActBCs, which, in addition to low CD21 expression, can be characterized by high CD71 expression, are thought to result from the GC pathway. ActBCs have been described as B cells that recently responded to their cognate antigens, as observed in influenza HA-vaccination (*16*, *17*) and more recently in COVID-19 (*18*). ActBCs peak in the circulation shortly after vaccination or infection and gradually decline over time with an accompanying increase in the frequency of MBCs, indicating a progressive differentiation of ActBCs into MBCs over time. ActBCs display an intermediate phenotype between MBCs and ASC and have also been suggested to be precursors of long-lived ASCs (*17*).

In SARS-CoV-2 infection, while ActBCs are gradually declining over time, affinity matured MBCs are accumulating. Remarkably, ActBCs are still detectable 6 month post-infection and during this period the overall BCR mutation count increases, supporting the notion of a persistent ongoing GC reaction in SARS-CoV-2 infection (*18*). Moreover, it has been suggested that patients with initial severe disease exhibit an impaired GC reaction, although severe COVID-19 is associated with persistent immune activation resulting in a stronger antibody response (*19*).

In this study, our goal was to investigate the phenotypic heterogeneity of circulating B cells with various antigen specificities in individuals with laboratory-confirmed SARS-CoV-2 infection. We studied individuals that exhibited a wide range of initial disease severity, and examined their B cell populations approximately 3-4 months after the onset of illness. At this time the GC reaction of SARS-CoV-2 specific B cells is still ongoing while the MBC compartment and ASCs have already started establishing for some time, providing a high-resolution snapshot of B cell and ASC lineages. In these samples, we simultaneously examined SARS-CoV-2 reactive B cells (Spike (S), receptor binding domain (RBD) and Nucleocapsid (NC)) generated in response to recent infection and MBCs that had been established prior to the SARS-CoV-2 pandemic (Influenza-HA-, RSV-F-, Tetanus-Toxoid-specific B cells). This analysis was conducted in a period where re-exposure to Influenza, respiratory syncytial virus (RSV) and travel was almost absent (summer 2020, shortly after stringent lockdown measures were gradually lifted in the Netherlands).

Unsupervised analysis of the B cell heterogeneity, as determined by high-dimensional flow cytometry (using 24 B cell markers), identified unique combinations of markers that distinguished novel B cell populations. These populations were then assessed in terms of disease severities, period of antigen exposure, and their phenotypic relation to other B cell and ASC subsets. This approach identified B cell subsets and novel markers characterizing active B cells as intermediates for generating ASCs or activated MBCs. The activated MBCs further contribute to the pool of resting long-lived MBCs.

## RESULTS

### Convalescent COVID-19 study cohort

To characterize the establishment and maintenance of B cell responses, we compared B cell responses against SARS-CoV-2 antigens in recent COVID-19 infection to those elicited by antigens of pathogens encountered in the past. PBMC samples for this study were collected from individuals who had experienced COVID-19 and were part of two prospective cohorts (*20*). We included individuals who had experienced various degrees of initial COVID disease, mild (N=33), moderate (N=18), severe (N=12), and critically ill COVID-19 disease (N=15). We selected samples 3-4 months post-onset of illness, at a time when GC reactions were still ongoing, but MBC and ASC formation had already been established (*19*, *21–23*). Each group was matched as much as possible for the age, gender, and collection time. Age ranged from 31 to 75 years old (median of 56, IQR 50-62) and did not significantly differ between the four groups of disease severity (Fig. S1A). The cohort consisted of 25 females and 53 males with men more likely to be in the group with critical COVID-19 disease (Fig. S1A). Blood samples were collected between 75 and 143 days after symptom onset (median of 98 days, IQR of 83-105 days), sample selection occurred slightly earlier for the mild COVID-19 group (median of 80 days IQR 77-89 days) compared to the other three groups (median of 103 days and IQR of 99-107 days). Additionally, a healthy donor control group (HC, N=11) that did not report COVID-19 infection and showed negative antibody titers to S and NC proteins was included in the present study.

### Antigen-specific B cell frequency is associated with disease severity and time since antigen exposure

First, the SARS-CoV-2-specific B cell response was examined for reactivity against S, RBD and NC across the range of COVID-19 disease severities. To do so, we developed a 29-color flow cytometry panel, comprising five channels for antigens detection, one Live/Dead stain, and the remaining 23 channels to detect surface antigens based on recent studies investigating the heterogeneity within human B cell compartments (Table S1) (*9*, *16*, *24*). Moreover, a combinatorial B cell staining approach with dual labeled antigenic probes was used, enabling simultaneous identification of B cells that are reactive to one of six different antigens in a single sample (*25*). In addition to the three SARS-CoV-2 antigen specificities that capture recently induced, antigen-driven B cell responses, B cells reactive to hemagglutinin (HA) from H1N1/pdm2009 influenza virus, fusion glycoprotein (F) from RSV and tetanus toxoid (TT) vaccine, were evaluated as representatives for B cell responses to pathogens or vaccines encountered in the past (Fig. 1A, S1B). Of note, during the period of sample collection (Summer 2020, shortly after the first wave of infections), influenza or RSV transmission were absent and travel was still strongly restricted, making DTP vaccination in these adult groups an unlikely event.

**FIGURE 1:**
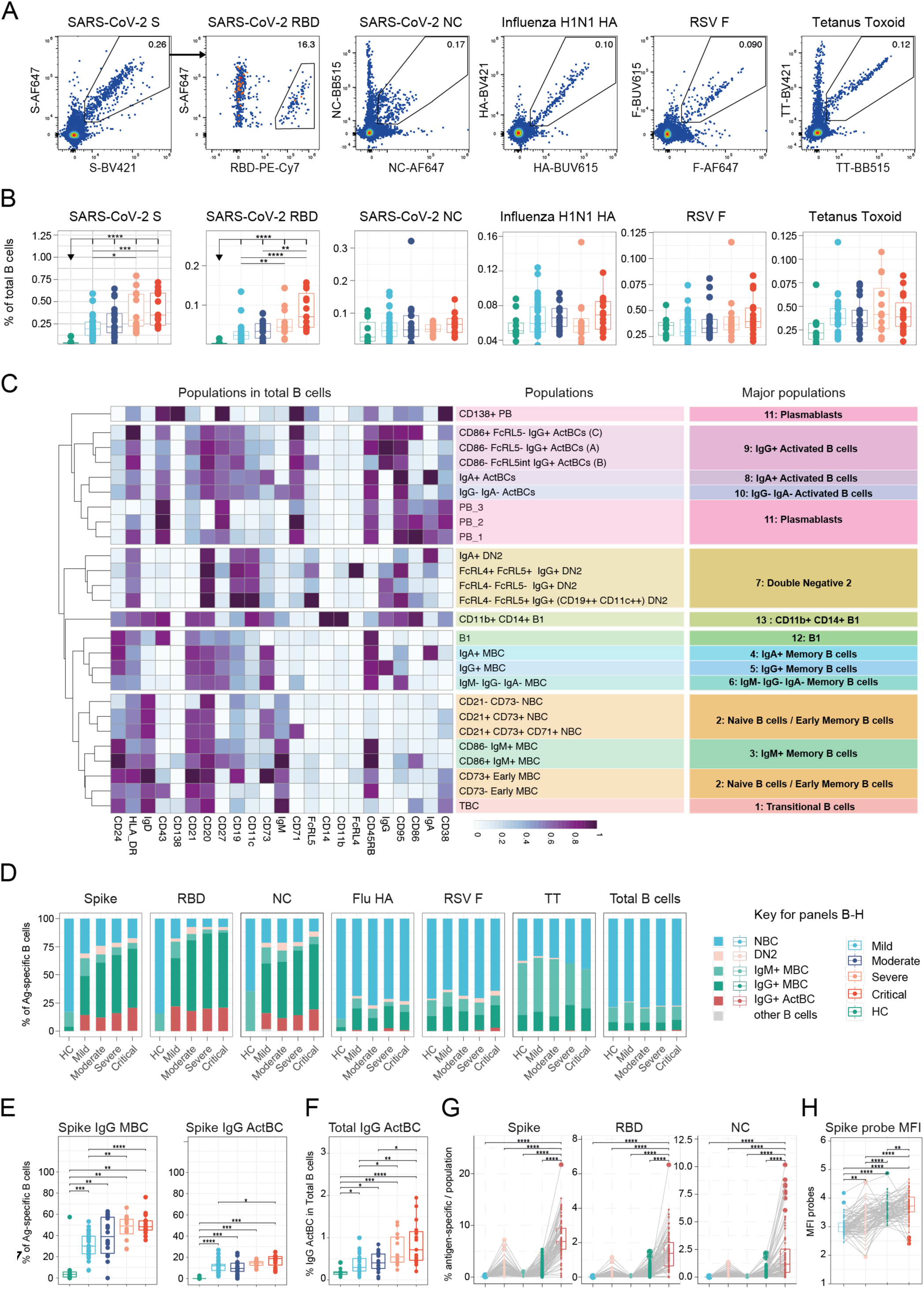
IgG+ Activated B cells and Memory B cells are engaged in SARS-CoV-2 response. **(A)** Combinatorial probe staining and gating strategy for the detection of multiple B cell specificities in a single PBMC sample. From live B cells (gating strategy Fig. S1B), antigen-reactive B cells are detected as double positive for the binding of the same antigen multimerized with two different fluorochromes. RBD specific B cells are detected out of S-specific B cells. **(B)** Frequency of antigen-reactive B cells in total B cells from Healthy controls, mild, moderate, severe and critical patients. **(C)** Heatmap of the 26 B cell populations and 13 Major populations defined after FlowSOM analysis of 23 parameters from total B cells of healthy controls and patients. **(D)** Frequency of B cell subsets defined by FlowSOM analysis according to B cell specificity and disease severity. **(E)** Frequency of IgG+ memory B cells (left) or IgG+ Activated B cells (right) in S-specific compartment according to disease severity. **(F)** Frequency of IgG+ Activated B cells in total B cells according disease severity. **(G)** Frequency of SARS-CoV-2 specific B cells (S: Left, RBD: middle, NC: right) in naive, DN2, IgM+ memory B cells, IgG+ memory B cells and IgG+ Activated B cells. **(H)** S probe Median fluorescence intensity in naive, DN2, IgM+ memory B cells, IgG+ memory B cells and IgG+ Activated B cells.

Frequencies of both S- and RBD-specific B cells within the total B cell compartment were significantly higher in each of the four COVID-19 severity groups compared to the HC group (%S/%RBD B cell specificity; HC: 0.034%/0.003%, mild: 0.193%/0.023%, moderate: 0.216%/0.033%, severe: 0.291%/0.044%, critical 0.349%/0.070%) (Fig. 1B). Furthermore, convalescent patients with severe or critical initial COVID-19 disease exhibited significantly higher SARS-CoV-2 S- and RBD-specific B cell frequencies compared to patients with mild disease. This heightened B cell response is associated with an increased antibody binding titers to S and RBD, along with augmented neutralization titers, as previously reported in these patients (*18*, *22*, *26*) (Fig. 1B, S1C). In contrast, the frequencies of B cells with specificity to SARS-CoV-2 NC and the three non-SARS-CoV-2 antigens did not differ between the four severity groups or between convalescent COVID-19 patients and the HC group.

Moreover, the frequency of SARS-CoV-2 S-specific B cells in patients was significantly increased in comparison to the three non-SARS-CoV-2 antigens (S: 0.233%, NC: 0.049%, HA: 0.062%, TT: 0.037%, RSV-F: 0.034%) consistent with the kinetics of B cell responses in circulation following recent infection and the ongoing GC activity at this time point (Fig. S1D) (*19*).

Taken together, these results demonstrate that the abundance of B cells reactive to a particular antigen in the circulation is influenced by the antigen properties (both qualitative and quantitative) and/or the time elapsed since antigen encounter, and the experienced severity of disease.

### Phenotypic heterogeneity of B cells is defined by antigen properties and time since exposure

To capture the heterogeneity of the circulatory B cell compartment, an unsupervised high-dimensional analysis of 23 surface antigens acquired by a spectral flow cytometer was used to define the main B cell subsets. This analysis led to the identification of 13 major populations aligning with the main circulating subsets described in the literature (*9*, *27*), which were further subdivided into 26 distinct subpopulations (Table S2) (Fig. 1C, S2A-B). These 13 major populations included unswitched B cells, comprising transitional B cells[1], naïve/early MBCs[2], and IgM MBCs[3]. Additionally, classical switched MBCs were identified, which were present in various isotypes (IgA+[4], IgG+[5], IgG-IgA-[6]). Responsive B cells, such as atypical double negative 2 (DN2) B cells[7] and Activated B cells (ActBCs) with diverse isotype profile (IgA+[8], IgG+[9], IgG-IgA-[10]), were also captured. This analysis further encompassed plasmablasts[11], defined as antibody-producing cells. Furthermore, rare innate B cells specifically B1 cells[12] and CD11b+ CD14+ B1 cells[13] were identified.

Subsequently, we compared the composition of B cell compartments specific to the six antigens. Our analysis revealed that five out of 13 major populations—NBC, DN2, IgM+ MBC, IgG+ MBC and IgG+ ActBC— accounted for most of the heterogeneity within the antigen-specific compartments (Fig. 1D). Notably, B cells specific for SARS-CoV-2 antigens in convalescent patients displayed a phenotype indicative of an ongoing response in comparison to B cells recognizing antigens encountered in the past and in relation to total B cells (Fig. 1D, S3A). Specifically, the IgG+ MBC emerged as the predominant subset within B cells reactive to SARS-CoV-2 proteins (IgG+ MBC; S: 40%, NC: 34%) surpassing other antigens (IgG+ MBC; HA: 14%, RSV-F: 16%, TT: 15%). Interestingly, 13% of S-specific B cells showed an ActBC phenotype (17% and 12% of RBD and NC-specific B cells respectively), a subset conspicuously absent in B cells specific to other antigens (IgG+ ActBC, HA: 1%, RSV-F: 0%, TT: 0%).

Furthermore, while the DN2 subset constituted a minor proportion of SARS-CoV-2 reactive B cells, it was significantly higher in comparison to other antigens (DN2, S: 4%, NC: 4% versus HA: 2%, RSV-F: 3%, TT: 1%, total: 1%). Specifically, DN2 B cells with specificity to S were highly enriched in FCRL5+ FCRL4+ IgG+ DN2 subset (DN2 FCRL5+ FCRL4+ IgG+ DN2, S: 30% vs total B cells: 10%; Fig. 1C, S3C) and were virtually absent from B cells with non-SARS-CoV-2 specificities (Fig. S3D).

Initial COVID-19 disease severity also impacted SARS-CoV-2 specific B cell subset distribution, as the IgG+ MBC subset significantly increased in abundance with disease severity (i.e. S-specific IgG+ MBC; Mild:30%, Moderate:39%, Severe:49%, Critical:48%) at the expense of the NBC and IgM+ MBC subsets. A similar trend was found for the ActBC subset, but was only statistically significant for the S specificity (i.e. S-specific IgG+ ActBC; Mild:12%, Moderate:9%, Severe:15%, Critical:19%) (Fig. 1D-1E, S3B). Of note, in the total B cell compartment, frequencies of S- and RBD-specific B cells with either DN2 (specifically FCRL5+ FCRL4+ IgG+ DN2 phenotype), IgG+ MBC or ActBC phenotype increased with disease severity (Fig. S3D-E). Thus, these three subsets account for the overall increase in S-specific B cells in severe and critical cases (Fig. 1B). Remarkably, the rise of the frequency of ActBCs with disease severity can also be observed in the total B cell compartment, even without gating on antigen-specific B cells. While low in frequency in the HC group, ActBCs are significantly more frequent in COVID-19 convalescent patients after 3-4 months following illness onsets (Fig. 1F, S3B), and can reach frequencies that exceed 1% in certain individuals who had severe or critical COVID-19 disease. This scarce ActBC population exhibited the highest enrichment in specificity to the recently encountered SARS-CoV-2 proteins (S: 6.9%, RBD: 1.4%, NC: 1.2%) (Fig. 1G), and likely represents recent emigrates released into the circulation from ongoing GC reactions (*17*). In addition, S-specific B cells with an ActBC phenotype displayed higher median fluorescence intensity (MFI) of the two fluorescent S-probes compared to the four other B cell subsets (Fig. 1H, S3F), suggesting a significantly increased avidity of the BCR on the surface of ActBCs, likely due to a prolonged affinity maturation (*28*).

These findings indicate that even several months after COVID-19 illness onset, the B cell response remains active. While antigen-specific ActBCs remain detectable in the circulation, likely resulting from ongoing GC reactions within lymph nodes, classical antigen-specific MBCs are emerging as well. This process appears to be more pronounced among individuals who experienced more severe COVID-19 disease. In stark contrast, B cells specific to antigens encountered prior to the onset of COVID pandemic are markedly less frequent in circulation and exhibit a quiescent phenotype. These observations highlight the dynamic nature of antigen-specific B cell responses.

### Comparison of ongoing and pre-pandemic antigen-specific B cell responses identifies active and resting memory B cell compartments

The strong enrichment of IgG+ ActBCs and IgG+ MBCs with specificity to SARS-CoV-2 indicate a recent reactivity of these two subsets to the viral infection. However, ongoing debate exists regarding the origins and outcomes of these subsets. Some hypotheses suggest that circulating ActBCs may precede the formation of MBCs, whereas others propose their involvement in the generation of ASCs (*16–18*, *29*). In our study, IgG+ ActBCs exhibited an intermediate profile between IgG+ MBCs and PBs, supporting both hypotheses (Fig. 1C and S3G). To gain a deeper understanding of the dynamics and heterogeneity between ActBCs and MBCs, a more exhaustive analysis was conducted, focusing on those cells with specificity to the six antigens.

Using dimensionality reduction and clustering techniques with highly variable surface proteins (listed in Table S3) as distinguishing features, four discrete clusters were identified among antigen-specific IgG+ MBCs and ActBCs (Fig. 2A-2B). Cluster 2 closely resembled the previously identified IgG+ ActBC subsets through FlowSOM clustering, while the remaining clusters (1, 3, and 4) predominantly corresponded to the IgG+ MBC subset (Fig. 2B-2C). B cells in Cluster 2 (IgG+ ActBCs) were restricted to SARS-CoV-2 antigen specificities (Fig. 2A-B, 2D). Additionally, B cells in cluster 2 displayed higher expression levels of CD38, CD43, CD71, CD86, HLA-DR, CD20, CD95, CD11c, and FcRL5, along with lower expression levels of CD24, and CD21, when compared to cells in the other clusters (Fig. 2E, S4A-B).

**FIGURE 2:**
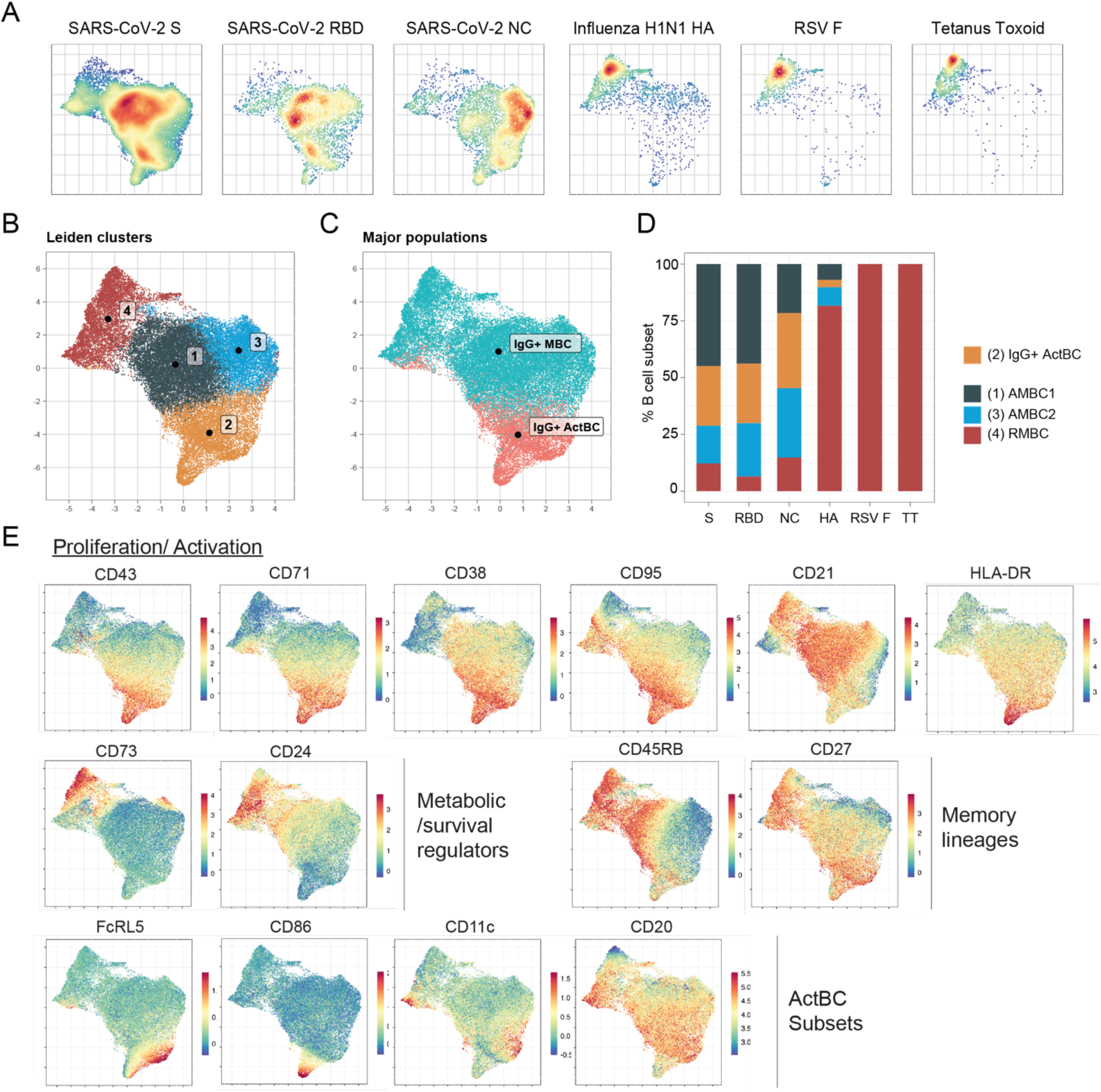
CD73 and CD24 separate true resting memory to activated memory B cells. **(A)** UMAP of flow cytometry data (13 markers, see table S3A) from all antigen-specific B cells captured from IgG+ memory B cells and IgG+ Activated B cells. UMAP representation of each B cell specificity (from left to right: S, RBD, NC, HA, RSV F, TT). **(B)** Leiden clustering identified 4 distinct clusters (1: Activated MBC1, 2: Activated B cells, 3: Activated MBC2, 4: Resting MBC) **(C)** Overlay of IgG+ memory and activated major populations captured by FlowSOM clustering on the UMAP data generated out of antigen-specific B cells. **(D)** Frequency of the 4 clusters as identified by Leiden clustering (1:Activated MBC1 , 2: Activated B cells, 3: Activated MBC2, 4: Resting MBC) according to antigen-specificity. **(E)** Feature plots showing scaled normalized counts for 14 relevant B cells markers in all selected cells.

Among the three IgG+ MBC populations (clusters 1, 3, 4), cluster 4 formed a distinct island from clusters 1 and 3 on the UMAP and was restricted to antigens encountered in the past. In contrast, B cells within clusters 1 and 3 were highly enriched in specificity to SARS-CoV-2 antigens and were more proximal to ActBCs (Fig. 2A-D). Of note, within SARS-CoV-2-specific IgG+ MBCs, the frequency of cluster 1 and 3 — but not cluster 4 — increased with disease severity and best correlated with antibody binding and neutralization titers (Fig. S4C-E).

Phenotypically, cluster 4 displayed more quiescent features, as evidenced by low levels of proliferation or activation markers (i.e. CD71, CD38, and CD43, HLA-DR), coupled with high levels of CD73, CD24 and CD45RB expression (Fig. 2E, S4A-B). Clusters 1 and 3, on the other hand, exhibited an intermediate phenotype, in between clusters 4 and 2, exhibiting moderate expression for markers CD71, CD43, CD38, HLA-DR and CD24, and predominantly negative for CD73 (Fig. 2E, S4A-B). Intriguingly, between clusters 1 and 3, cells in cluster 1 were enriched in specificity to SARS-CoV-2 S and RBD antigens, while cells in cluster 3 were enriched in specificity for NC (Fig. 2A-D, S4F). These two clusters were distinguishable phenotypically; cluster 1 displayed higher levels of CD24, CD21, CD27, CD45RB, and CD95 expression, whereas cluster 3 exhibited a CD45RB− CD27^low^ CD21^+^ CD11c^−^ phenotype (Fig. 2E, S4A-B). This latter profile bears resemblance to the previously described double-negative 1 (DN1) B cells (*11*, *30*).

Based on these results, we propose that cluster 4 represents the “true” resting MBC (RMBC) compartment, whereas clusters 1 and 3 represent recently activated MBC (AMBC1 and AMBC2, respectively). Our analysis of differential phenotypic expression suggests a progressive transition from ActBCs to RMBCs, featuring an intermediate phase represented by AMBCs. The transition from AMBCs to RMBCs is marked by the concomitant acquisition of CD73 and CD24. These markers have previously been implicated in metabolic regulation and survival of B cells, respectively (*31*, *32*).

### The resting memory B cell compartment is heterogenous and contains long-lived memory B cells

The RMBCs within the IgG+ MBC compartment demonstrated significant phenotypic heterogeneity for specific surface proteins, including CD73, CD95, CD24, CD45RB, CD38 and CD21 (cluster 4, Fig. 2B, 2E, S4B). This heterogeneity was not captured by the previous analysis of antigen-specific IgG+ MBCs and ActBCs (Fig 1C, 2B, 2E). To refine our understanding of the RBMC compartment, we expanded the population to include IgG+ MBCs that were unspecific to any of the six antigens into the pool of antigen-specific IgG+ MBCs.

Dimensionality reduction and clustering analysis of this new composite dataset corroborated the distinct identification of RMBCs (clusters 1, 5, 6, 7, 8, 9) apart from AMBCs (clusters 2, 3, 4) (Fig. 3A, S5A). As expected, SARS-CoV-2 specificity was predominantly confined to AMBCs, whereas the specificity to other antigens and non-specific IgG+ MBCs were mainly present in RMBCs (Fig. 3A-3C).

**FIGURE 3:**
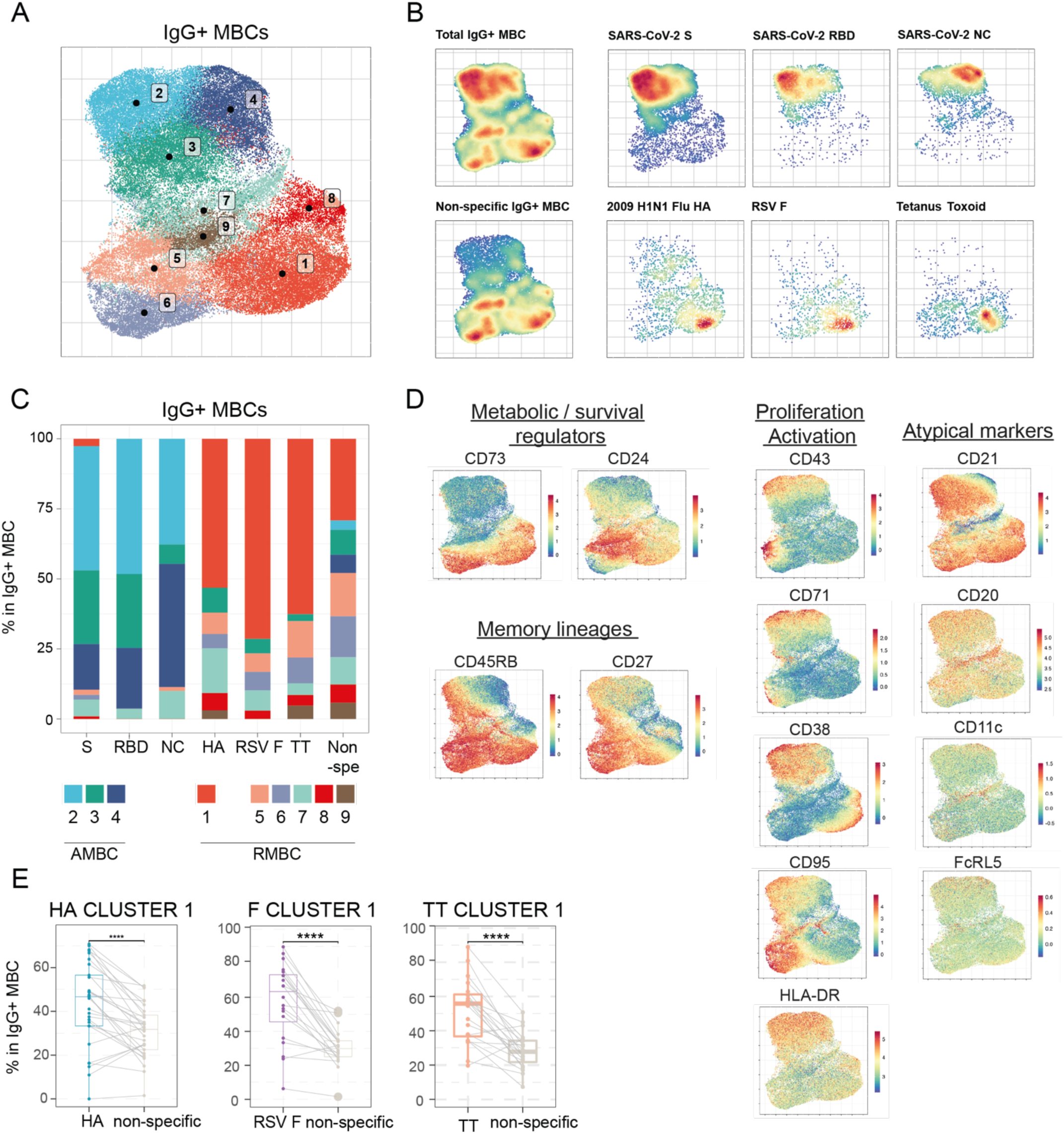
Resting memory B cells encompass multiple subsets that can be segregated based on CD73, CD24, and CD95 expression. **(A-E)** Flow cytometry input data originates from all antigen-specific in addition to 1000 non-specific B cells from each individual donor of the IgG+ memory B cells major population defined by FlowSOM clustering. **(A)** Leiden clustering (14 markers, see table S3B), nine clusters were identified. **(B)** UMAP of the selected flow cytometry data. UMAP representation of each B cell specificity (from left to right: total selected data, S, RBD, NC, HA, RSV-F, TT, non-specific B cells). **(C)** Frequency distribution of the nine clusters as identified by Leiden clustering according to antigen-specificity. Only data points corresponding to a minimum of 20 antigen specific B cells were used for the analysis. **(D)** Feature plots showing scaled normalized counts for 13 relevant B cells markers in all selected cells. **(E)** Comparative analysis of cluster 1 for HA, RSV F, and TT versus non-specific B cells, for samples that encompass at least 20 cells for a given specificity.

Six distinct clusters can now be defined within the RMBC population. Among these, cluster 8 lacked CD45RB expression and exhibited a resting DN1-like phenotype (CD27low CD45RB− CD21+ CD11c−). Cluster 7 displayed a significantly reduced level of CD21 expression and demonstrated notable heterogeneity with respect to other markers, such as CD11c and FcRL5 that may describe other atypical B cell subsets (*33–35*)(Fig. 3A, 3D, S5B).

The remaining RMBC clusters (1, 5, 6, 9) were characterized as CD45RB+ CD21+ and could be further subdivided into two distinct groups based on CD95 expression: CD95+ clusters (clusters 5 and 6) and CD95− clusters (clusters 1 and 9). While CD95+ clusters showed some degree of activation, as indicated by the expression of CD71 and CD43, the CD95− clusters displayed a more resting phenotype. Moreover, expression levels of CD24 and CD73 further subdivided both CD95+ and CD95− clusters. Indeed, CD95+ populations consisted of CD24hi CD73−/+ (cluster 5) and CD24lo CD73hi (cluster 6) subsets. Similarly, CD95− cluster 1 displayed high levels of CD73 while expressing moderate levels of CD24, and was also distinguishable in the RMBC compartment by its expression of CD38. Cluster 9, on the other hand, showed a CD24hi CD73− phenotype within the CD95− subset (Fig. 3A, 3D, S5B, S5C). Overall, RMBCs with high CD73 expression tended to exhibit moderate CD24 levels, while the converse was also true; the high expression of one of these makers is unique to the RMBC compartment (Fig. 3D, S5B, S5C).

Investigation of cluster distribution within the RMBC compartment with respect to antigen specificities, revealed that B cells lacking antigen specificity were strongly dominated by three clusters (cluster 1: 28%, cluster 5: 15%, cluster 6: 14%). In contrast, a pronounced bias to cluster 1 (50-60%) was found among B cells with specificity to TT, RSV-F, and HA antigens (Fig. 3E, S5D). Of note, cluster 8 and 9 had limited representation (<2.5%) over all B cell specificities.

In conclusion, the RMBC population is distinguishable by high levels of CD24 and/or CD73 expression and can be subdivided into three subsets: two CD95+ subsets displaying residual signs of activation or proliferation, and a quiescent CD95− CD73+ CD24lo MBC population. Notably, the CD95− CD73+ CD24lo subset is particularly enriched with B cells responsive to antigens encountered years or even decades prior, indicating a long-lasting persistence.

### Activated B cells are poised for distinct lineages

Given that ActBCs have been identified as potential precursors of ASCs (*17*, *27*), and that both ActBCs and ASCs can be further subdivided into multiple subsets using FlowSOM clustering, our subsequent analysis aimed to unravel the phenotypic relationship between these sub-populations. Briefly, FlowSOM clustering identified 4 distinct populations that were characterized as ASCs (Fig. 1C); a population of plasmablasts precursors (PB_1; CD19int, CD20int, CD27+ CD38+ CD138−, 0.055%) and two populations of immature plasmablasts (PB_2 and 3; CD19lo, CD20−, CD27hi CD38hi+ CD138−, 0.086% and 0.062%) and a small population of mature CD138+ plasmablasts (CD19low, CD20low, CD27hi CD38hi CD138+, 0.003%) (*36*, *37*). Furthermore, within the IgG+ ActBC compartment, FlowSOM clustering identified 3 populations, driven principally by the expression of FcRL5 and CD86; ActBC A (FcRL5− CD86−, 0.308%), ActBC B (FcRL5+ CD86−, 0.073%), and ActBC C (FcRL5+ CD86+, 0.046%) populations. Of interest, we observed an increase in ASC frequency correlating with heightened disease severity, accompanied by a shift from pre-plasmablast B to immature plamablast dominance (Fig. S6A). However, disease severity did not affect the distribution of ActBCs; instead, antigen specificities played a significant role, with B cells specific to the NC protein exhibiting an increase in ActBC A frequencies and a decrease in ActBC B frequencies compared to those specific to the S protein (Fig. 4A, S6B).

**FIGURE 4:**
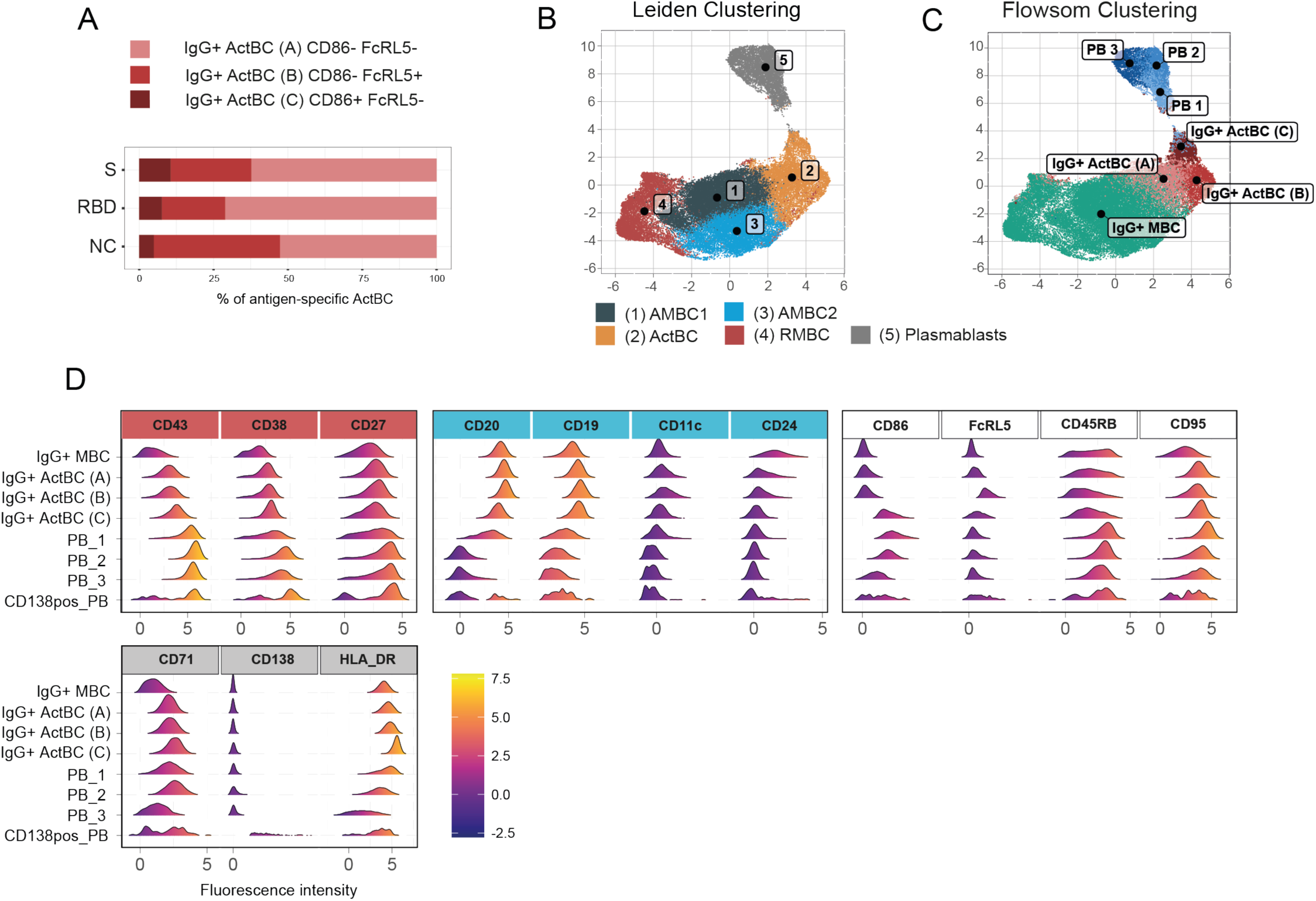
Activated B cells at the crossroad of Memory and antibody-secreting cells. **(A)** Frequency of IgG+ Activated B cell populations (A: CD86− FcRL5−, B: CD86− FcRL5+, C: CD86+ FcRL5−) out of SARS-CoV-2-specific B cells (S, RBD, NC). **(B-C)** UMAP analysis of flow cytometry (15 markers, see table S3C) generated out of all plasmablasts and all antigen specific B cells captured from IgG+ memory B cells and IgG+ Activated B cells. **(B)** Overlay of plasmablasts, clusters 1-4 (previously generated in figure 2 as identified by Leiden clustering out of IgG+ Activated B cells and memory B cells), on the UMAP **(C)** Overlay of activated populations (A: CD86− FcRL5−, B: CD86− FcRL5+, C: CD86+ FcRL5−), plasmablasts population, and IgG+ MBCS, all captured by FlowSOM clustering on the UMAP data **(D)** Comparative analysis of cell surface expression by histogram representation of 14 relevant B cells markers between populations of IgG+ Activated B cells (A: CD86− FcRL5−, B: CD86− FcRL5+, C: CD86+ FcRL5−) and plasmablasts (PB_1, PB_2, PB_3, CD138pos_PB).

To investigate the relationships among these populations, we constructed a composite dataset that encompassed ASCs and antigen-specific IgG+ ActBCs and IgG+ MBCs. Dimensionality reduction and clustering analysis revealed a phenotypic continuum from IgG+ ActBCs to ASCs, with the IgG+ ActBC population C exhibiting the closest phenotypic resemblance to the pre-plasmablast PB_1 (Fig. 4B-D, S6C-D). This population C lies at the intersection of ActBCs and ASCs. Along this continuum, we observed an upregulation in expression of ASC lineage markers, including CD43, CD27, and CD38, accompanied by a downregulation of CD20, CD19, CD11c, and CD24. Notably, CD86 exhibited transient expression in ActBC population C, PB_1, and PB_2, but this expression was diminished in mature plasmablasts. On the other hand, both ActBC A and B were phenotypically closer to the AMBC populations. Consequently, it is tempting to speculate that IgG+ ActBCs in population C (CD86+) are predisposed to differentiate into ASCs, while IgG+ ActBCs in populations A and B (CD86−) are predisposed to differentiate into MBCs. Compared to ActBC A, population ActBC B was positive for FcRL5, mostly CD45RB- and displayed a lower expression of CD95 (Fig. 4D, S6C-D). Interestingly, both ActBC A and AMBC2 displayed a CD45RB− and CD95lo phenotype and were enriched in NC specificity, implying a potential relationship between these two populations (Fig. 2, 4A, S6D).

Collectively, these findings show that blood IgG+ ActBCs are enriched in specificity to recently encountered antigens and are phenotypically heterogeneous. Moreover, distinct phenotypes might define populations with different developmental trajectories and fates.

## DISCUSSION

This study sought to elucidate human B cells differentiation by investigating the composition of the B cell compartments of various antigen specificity at different times of exposure. Specifically, we aimed to identify distinctive cell surface protein signatures that characterize undescribed and meaningful subsets of experienced B cells and precursors of long-lived MBCs and ASCs. Using high-dimensional flow cytometry and unsupervised analysis, we compared B cells from convalescent COVID-19 patients, with specificity to antigens encountered prior to the pandemic (HA, RSV-F, and TT) and B cells with specificity to recently encountered SARS-CoV-2 antigens (S, RBD, and NC), across varying COVID-19 disease severities. This analysis enabled the identification of distinct B cell phenotypes that give rise to MBCs and ASCs, offering a novel panel of cell surface proteins to distinguish between Activated B cells (ActBC), ASC lineage, activated MBCs (AMBC) under selection, “true” resting MBCs (RMBC), and long-lived MBCs.

Using this approach, we first identified two notorious subsets: IgG+ ActBCs, limited to B cells with specificities to SARS-CoV-2, and IgG+ MBCs which exhibited a dominant response to SARS-CoV-2. The origin and fate of ActBCs has remained a subject of debate. Here, we demonstrated that the expansion of these subsets correlated with disease severity and were not present in the bloodstream of seronegative healthy controls. Additionally, SARS-CoV-2 specific ActBCs had higher relative affinity to S protein compared to other B cells and correlated with both SARS-CoV-2 IgG and neutralization titers. This suggests that 3-4 months after illness onset, ActBCs arise from GC reaction, as it has been proposed by other groups (*17*).

Previous studies have shown the progressive differentiation of ActBCs into MBCs using B cell repertoire analysis (*16*, *18*). Yet, our data suggest the potential of a phenotypic transition from ActBCs to MBCs as well. We delineated three undescribed IgG+ MBC populations: a “true” RMBC population restricted to antigen encountered in the past, and two AMBCs populations that were restricted to SARS-CoV-2 antigen specificities and clustered in close proximity to ActBCs. Our findings indicate a phenotypic continuum from ActBCs to RMBCs through the intermediary AMBC population, with the acquisition of CD24 and CD73 distinguishing AMBCs from RMBCs. Importantly, CD21+ B cells, which have long been characterized as resting MBCs, are actually a mix of AMBC and “true” RMBCs. Three to four month post-COVID infection, nearly all CD21+ B cells with SARS-CoV-2 specificities show residual activation and belong to the AMBC CD21+ CD73-CD24-subset.

Moreover, within the RBMC compartment, CD24 and CD73 were conversely expressed with a gradient transitioning from CD73hi to CD24hi. In tandem with the CD95 marker, they divided the RMBC compartment into three distinct populations, including two CD95+ B cell populations that exhibit residual proliferative and activated states, and a quiescent CD95− CD73+ CD24lo population highly enriched in B cell reactive to antigens encountered longer ago. Our findings introduce this CD95− CD73+ CD24lo population as a defining phenotype of IgG+ circulating long-lived MBCs in humans.

CD95 or Fas is known as a death receptor that triggers apoptosis in response to FasL interaction in absence of survival signals (*38*). This mechanism is potentially pivotal for the selection of CD95− long-lived MBCs, particularly when post-infection inflammation subsides and antigen availability decreases. On the other hand, CD24 and CD73 likely contribute to the establishment of the MBC population. Their roles in establishing quiescent naive follicle B cells (*31*) suggest parallels in MBC development. Indeed, B cell selection in the bone marrow involves CD24 and BCR specificity, leading to apoptosis of autoreactive B cells (*39*, *40*) culminating in the emergence of transitional B cells, expressing high CD24 levels, in the periphery. Our findings indicate a gradual loss of CD71 and gain of CD24 expression along the B cell differentiation trajectory to RMBCs, emphasizing a critical role for CD24 in the selection or development of resting memory cells. Next, as transitional B cells differentiate into follicular B cells, CD73 expression increases while CD24 decreases (*31*). Functionally, CD73, in tandem with CD39, converts ATP to immunosuppressive adenosine — termed the adenosine salvage pathway — fostering metabolic quiescence in naive B cells. In the RMBC compartment, CD24 and CD73 exhibit a similar inverse gradient of expression, defining various B cell populations. We have shown that CD95− CD73+ CD24lo expressing RMBCs are preferentially enriched in the long-lived memory cells. These cells may exhibit metabolic quiescence, akin to naive B cells (*31*) and CD73+ memory T cells (*41*, *42*), that can lead to long-lived properties. Additionally, CD95− CD73+ CD24lo B cells are the only cells within RMBCs that express CD38, pointing towards non-canonical adenosinergic pathways that are CD39-independent (*43*). Supporting the importance of CD73 in B cell survival, recent work identifies CD73+ subset as antigen-experienced B cells (*9*) that is preferentially expressed in isotype-switched B cell population. Additionally, long-lived splenic anti-smallpox MBCs were restricted to a CD27+ CD21+ CD73+ phenotype and displayed long-lasting GC imprinting (*44*). Our study establishes CD73 and CD24 as reliable markers to distinguish RMBCs from AMBCs and underscores the selective nature of long-lived MBC formation.

In addition to their memory precursor competence, ActBCs has also been described as potential precursors of long-lived ASCs (*17*, *29*). In this study, we confirm that ActBCs exhibit an intermediate phenotype at the crossroad between MBCs and circulating ASCs (plasmablasts). Distinctively, we find that ActBCs can be further separated into three subsets based on the expression of FcRL5 and CD86. Among these; CD86+ FcRL5− ActBCs were phenotypically most similar to pre-plasmablasts, hinting that they may be poised for ASC differentiation. Conversely, the two other ActBC (CD86−) subsets may predominantly contribute to the MBC pool.

In the GC, CD86+ B cells are abundant, especially in the light zone (*45*). The upregulation of CD86 on B cells is stimulated by interleukin-21 (IL-21) signaling originating from CD4+ T cells (*46*). In this context, CD86 plays a pivotal role as a costimulatory molecule, facilitating crucial interactions between GC B cells and Tfh cells via the CD28 pathway (*47*, *48*). Its expression on ActBCs might denote recent CD4+ T cell help, likely occurring during the GC reaction. This is further supported by the more pronounced activated profile of the CD86+ FcRL5− ActBC subset (CD38hi, CD71hi, CD43hi, HLA-DRhi). Stable interaction with Tfh cells via costimulatory molecules is crucial for GC B cell differentiation into plasma cells (*49*), suggesting that activated CD86+ B cells have the potential to differentiate into ASCs. The exact fate of circulating CD86+ ActBCs as plasma cells, plasmablasts, or MBCs requires further investigation.

Our results also show that the ActBC CD86− are closer phenotypically to AMBC, suggesting their potential as MBC precursors. Based on FcRL5 and CD45RB expressions, the CD86− ActBC population can be separated into ActBC A (FcRL5− CD45RB+) and ActBC B (FCRL5+ CD45RB−) subsets. FcRL5 is an IgG receptor with both activating and inhibitory functions (*50*). In the absence of CD21, FcRL5 preferentially inhibits B cells (*51*), and its expression is associated with exhausted functionalities in chronic infection or autoimmunity (*10*). This might suggest potential inhibitory features in CD21low ActBCs expressing FcRL5. Intriguingly, while the FcRL5− ActBC subset was enriched in specificity to the S, the FcRL5+ ActBC subset was enriched in specificity to the NC antigen. This discrepancy may be due to the fact that NC antigen is relatively more conserved in coronaviruses (*52*), principally resulting in repeated stimulation from recurring infections throughout life. Additionally, the context of antigen presentation may influence the fate of these B cells. FcRL5+ B cells might exclusively react to membrane-bound antigens (*53*, *54*), potentially causing NC-specific FcRL5+ B cells, which respond to soluble antigens, to remain unresponsive and accumulate as FcRL5+ ActBCs in the bloodstream. Recently, FcRL5+ B cells have been depicted as recent GC graduates in one study and ASC precursors in another (*55*, *56*), our results suggest that differentiation in ASC and classical MBCs is accompanied by the loss of FcRL5.

Interestingly, the AMBC compartment was also separated into two populations, AMBC1, enriched for NC with an activated DN1-like phenotype (CD45RB− CD27lo CD21+), and AMBC2, enriched for S antigens (CD45RB+ CD27+ CD21+). While DN1 B cells have not yet been associated with pathology or function, we highlight here that NC-specific B cells were already enriched in a CD86− FcRL5+ CD45RB− CD27+ phenotype within ActBCs, suggesting the possibility of a phenotypic imprinting of the CD45RB expression that could support the transition from ActBCs to AMBCs. Also, in comparison to the S antibody response, NC showed low antibody durability (*57*), which could be seen as evidence for recall rather than primary response that would trigger short lived plasmablast and/or an EF response. Therefore, further research should determine whether CD86− FcRL5+ ActBCs and activated DN1 subsets could be the cellular counterpart of such processes.

Overall, these results demonstrate that distinct antigens from a single pathogen can elicit different B cell responses, underscoring the impact of antigen attributes including the quantity, quality, localization, pre-existing response, and context of antigen presentation on the fate of B cell differentiation.

In conclusion, our results depict that IgG+ B cells can be classified into three main populations: (i) Activated B cells that have recently emerged from the germinal center reaction and have the potential to differentiate to antibody-secreting cells or memory B cells, depending on CD86 expression, (ii) Activated memory B cells that arose from Activated B cells, still displaying residual activation, but likely on trajectory to become resting memory B cells, (iii) resting memory B cells, which can be distinguished from the two other subsets by their converse CD24 and CD73 expression signature. Importantly, resting memory B cells displaying a CD95− CD73+ CD24lo phenotype are highly enriched in specificities that were generated years to decades ago. These markers have been linked to metabolic quiescence and survival, potentially contributing to their long-lasting properties.

Our discoveries may be utilized as a valuable reference for subsequent research endeavors exploring the humoral immune response following vaccination or disease. The monitoring of these B cell subsets is informative in evaluating the immune response’s quality and devising effective intervention strategies. Tracking Activated B cells over time as surrogates for germinal center activity can guide vaccination strategies, aiding in pinpoint the optimal stage for boosting the immune response. Additionally, assessing the antibody-secreting cells precursors and the development of long-lived memory B cells are crucial aspects in examining the potency and durability of humoral responses to vaccination. In the context of disease, identifying markers that distinguish active B cells (Activated B cells and Activated memory B cells) highly enriched in responsive B cells can aid in uncovering B cells with currently unknown specificities. This is crucial for tracking and studying autoreactive B cells during active autoimmune diseases, as well as active anti-tumor B cells and B cells responsive to novel pathogen encounters. Finally, the revealed sequence of B cells at various stages of IgG+ memory B cells and antibody-secreting cells formation brings us closer to understanding the critical decision point for the bifurcation of IgG+ memory B cells and antibody-secreting cells. This information is essential for identifying regulators that can either enhance antibody formation against pathogens or intervene in undesired antibody formation in allergy, auto-, and allo-immune responses.

## MATERIALS AND METHODS

### Human study design and clinical samples

Individuals with mild, moderate, severe and critical initial COVID-19 disease were followed in cohorts at Sanquin Blood Bank and Amsterdam UMC locatie AMC/Public Health Service of Amsterdam (*20*), Amsterdam, the Netherlands. Clinical severity was defined according to World Health Organization (WHO) criteria. The study sample was selected from these two cohorts so that individuals in the four groups of disease severity were matched by age and gender. Healthy controls blood was collected from healthy blood donors by a Dutch blood bank (Sanquin, Amsterdam).

### Study approval

Data and samples were collected only from voluntary, non-remunerated, adult donors who provided written informed consent as part of routine donor selection and blood collection procedures that were approved by the Ethics Advisory Council of Sanquin Blood Supply Foundation. Data and samples from Amsterdam UMC, location AMC, were collected only from voluntary, non-remunerated, adult individuals. Written informed consent was obtained from each study participant. The study design was approved by the local ethics committee of the Amsterdam UMC (Medisch Ethische Toetsingscommissie [METC]; NL73759.018.20) (*20*). Healthy controls donors consent was waived due to anonymized donation of blood for blood donation, blood products and research by the donors to the Dutch national blood bank. The study is in accordance with the declaration of Helsinki and according to Dutch regulations.

### Peripheral blood mononuclear cells isolation

Peripheral blood mononuclear cells (PBMC) were isolated from heparinized blood samples using standard Ficoll-Paque Plus gradient separation (GE Healthcare, Chicago, IL, USA). Cells were stored in 10% dimethyl sulfoxide in fetal bovine serum (Thermo Fisher Scientific, Waltham, MA, USA). Healthy and COVID-19 participant blood specimens were ficoll gradient separated into plasma and PBMCs. PBMCs were cryopreserved in FBS + 10% DMSO for future use.

### Protein design and purification

All soluble proteins, including SARS-CoV-2 S-2P (*58*), RBD, influenza A hemagglutinin (H1N1pdm2009, A/Netherlands/602/2009, GenBank: CY039527.2 (*59*)), RSV prefusion stabilized F (DS-Cav1 (*60*)), constructs with avi-tag and/or hexahistidine (his)-tag and/or strep-tag were expressed and purified as previously described ((*58*). After purification, avi-tagged proteins were biotinylated with a BirA500 biotin-ligase reaction kit according to the manufacturer’s instruction (Avidity). TT was purchased from Creative Biolabs (Vcar-Lsx003). NC and TT were aspecifically biotinylated using EZ-Link Sulfo-NHS-LC-Biotinylation Kit (Thermo Fisher) according to the manufacturer’s instructions.

### Probe preparation for detection of antigen-specific B cells

Biotinylated protein antigens were individually multimerized with fluorescently labeled streptavidin (BB515, BD Biosciences; BUV615, BD Biosciences; AF647, Biolegend; BV421, Biolegend) as described previously (*25*). Briefly, biotinylated proteins and fluorescently labeled streptavidin were mixed at a 2:1 protein to fluorochrome molar ratio and incubated at 4°C for 1 h. Unbound streptavidin conjugates were quenched with 10 uM biotin (Genecopoiea) for at least 10 min. A combinatorial probe staining strategy was used for simultaneous identification of multiple B cell specificities in a single sample. This combinatorial probe staining strategy uses all possible combinations of two fluorophores to increase the number of specificities that can be detected and decrease aspecific binding. In our study, we were able to detect 6 different antigen-specificities using 5 distinct fluorophores. Probes were labeled in the following manner: SARS-CoV-2 S (AF647, BV421), H1N1 HA (BUV615, BV421), RSV-F (AF647, BUV615), NC (AF647, BB515), TT (BB515, BV421), and RBD (PE-Cy7). Individual labeled proteins were then equimolarly mixed and kept at 4°C before usage. A final concentration of 45.5 nM of each probe is used to label B cells.

### Sample staining

The Antibody mix was supplemented at 10 uL with BD Horizon™ Brilliant Stain Buffer Plus (BD Biosciences, Franklin Lakes, NJ, USA) to minimize staining artifacts commonly observed when several BD Horizon Brilliant dyes are used. 10^7^ previously frozen PBMC samples were first depleted for T cells using CD3 selection kit II (StemCell) according to the manufacturer’s instruction. Enriched B cells were then stained at 4 °C for 30 mins with the mix of multimerized proteins and the mix of fluorochrome-conjugated antibodies simultaneously (Table S1). Following staining, cells were washed twice with a washing buffer containing 1% bovine serum albumin (Sigma-Aldrich, Saint Louis, USA) and 1 mM ethylenediaminetetraacetic acid (EDTA) in phosphate-buffered saline (Fresenius Kabi, ’s-Hertogenbosch, The Netherlands), then fixed with cold paraformaldehyde 1% for 10 mins at room temperature on a shaker and then washed twice with the washing buffer. Samples were acquired on a Cytek® Aurora 5 Laser (UV, V, B, YG, R) spectrum cytometer. Acquisition, spectral unmixing using reference controls, and analysis were performed using Cytek SpectroFlo® V3.0.1 software (Cytek Biosciences, Fremont, California, United States).

### Isotype-specific antibody ELISA

IgM, IgG and IgA to RBD and NCP were measured as described previously (Steenhuis et al., 2021). RBD and NP proteins were produced as described before (Steenhuis et al., 2021). Pooled convalescent plasma or serum was included on each plate as a calibrator (set to a value of 100 AU/mL) to quantify the signals. Results were expressed as arbitrary units (AU) per mL (AU mL) and represent a semi-quantitative measure of the concentrations of IgG, IgA and IgM antibodies to RBD and NP.

### Pseudovirus neutralization assay

Pseudovirus was produced by co-transfecting the pCR3 SARS-CoV-2–SΔ19 expression plasmid with the pHIV-1NL43 ΔEnv-NanoLuc reporter virus plasmid in HEK293T cells (American Type Culture Collection, CRL-11268) (*61*, *62*). Cell supernatant containing the pseudovirus was harvested 48 hours after transfection and stored at −80°C until further use.

HEK293T/ACE2 cells provided by P. Bieniasz (*61*) were seeded at a density of 20,000 cells per well in a 96-well plate coated with poly-l-lysine (50 μg/ml) 1 day before the start of the neutralization assay. NAbs (1 to 50 μg/ml) or heat-inactivated sera samples (1:100 dilution) were serial diluted in five fold resp. threefold steps in cell culture medium [Dulbecco’s modified Eagle’s medium (Gibco) supplemented with 10% fetal bovine serum, penicillin (100 U/ml), streptomycin (100 μg/ml), and GlutaMAX (Gibco)], mixed in a 1:1 ratio with pseudovirus, and incubated for 1 hour at 37°C. These mixtures were then added to the cells in a 1:1 ratio and incubated for 48 hours at 37°C, followed by a PBS wash, and lysis buffer was added. The luciferase activity in cell lysates was measured using the Nano-Glo Luciferase Assay System (Promega) and GloMax system (Turner BioSystems). Relative luminescence units were normalized to the positive control wells where cells were infected with pseudovirus in the absence of NAbs or sera. The inhibitory concentration (IC50) and neutralization titers (ID50) were determined as the NAb concentration and serum dilution at which infectivity was inhibited by 50%, respectively, using a nonlinear regression curve fit (GraphPad Prism software version 8.3) (45). Samples with ID50 titers of <100 were defined as having undetectable neutralization.

### Spectral flow cytometry data pre-processing

Using FlowJo, Flow Cytometry Standard (FCS) files were gated on single and viable cells, cells positive for CD3, CD4, CD16 or CD56 were excluded in a dump channel, and then CD19 was used to identify B cells (gating strategy is shown in Fig. S1). Antigen-specific B cells were then selected based on the combination of fluorochrome-conjugated streptavidin as is shown in Fig.1A. To remove potential cross-reactive B cells to streptavidin, each combination was first gated on cells that are double negative for the other two channels (Fig.S1).

Gated data was further processed with the R programming language (http://www.r-project.org) and Bioconductor (http://www.bioconductor.org) software. Initially, antigen-specificity was integrated as a logical variable in the data. Default setting of ’flow_auto_qc’ function from flowAI (Monaco et al., 2016) was used to detect and remove flow cytometry anomalies in both signal acquisition and dynamic range. Data was transformed with an inverse hyperbolic sine (asinh) transformation. Batch effects were modeled using reference samples stained and acquired with each batch to control for signal fluctuation that might occur over time due to changes in instrument performance. The model was then used to remove batch effects from the data using a normalization algorithm. Modeling of batch effect and data normalization was done using the CytoNorm package(*63*) in R.

### FlowSOM-based clustering

Following the data preprocessing step, we utilized FlowSOM for unsupervised clustering of the flow cytometry data. FlowSOM leverages a self-organizing map (SOM) algorithm and hierarchical consensus meta-clustering to cluster cells based on their phenotypic markers, enabling the identification of phenotypically defined populations (*64*, *65*). We included 23 surface proteins as input features. The FlowSOM algorithm was configured to use a self-organizing map (SOM) with a grid size of 20 × 20, resulting in 400 nodes. Nodes were then meta-clustered using the ’ConsensusClusterPlus’ function with k = 40 for hierarchical consensus clustering as implemented in the ConsensusClusterPlus package(*66*) in R, providing a comprehensive overview of distinct meta-clusters in total B cells.

### Heatmap visualization and met-cluster annotation

Heatmap was employed as a visual tool to interpret and illustrate the complex relationships inherent in FlowSOM-based meta-clusters. To that end, a random subset of the cells from the dataset were selected and grouped per meta-cluster and the median unscaled expression of the 23 surface proteins was computed. These median expression values were then used to construct a heatmap, visualizing the relationship between meta-clusters and cell surface protein expression using a hierarchical clustering dendrogram. The heatmap was generated using the ’make.pheatmap’ function from Spectre package(*67*) in R. Subsequently, the meta-clusters were annotated, and certain neighboring meta-clusters, which did not exhibit biologically significant differential expression of cell surface proteins, were merged for a more coherent representation.

### Dimensionality reduction

We used Uniform Manifold Approximation and Projection (UMAP) for non-linear dimensionality reduction of composite datasets using the ’run.umap’ function (neighbours = 15, min_dist = 0.1) from Spectre package in R. These datasets comprised a carefully curated selection of B cells, based on antigen specificities and distinct B cell subsets. We focused on IgG+ MBCs, IgG+ ActBCs and ASCs as identified by FlowSOM-based clustering and annotation. When required, we augmented the dataset of antigen-specific B cells by incorporating a randomly selected subset of antigen non-specific B cells. This addition was designed to ensure proportional representation of all subjects within the cohort to maintain the balance of our study population. Cell surface proteins were meticulously selected as features for dimensionality reduction based on its relevance to B cell activation, proliferation, antigen experience and metabolic regulation and on their expression variance within each composite dataset (Tables S3A-C). We employed an unsupervised clustering approach using community detection based on Leiden algorithm as embedded in ’cluster’ function from seqGlue package. Together, UMAP and the Leiden algorithms form a robust analytical framework for dissecting complex relationships and structures within the flow cytometry composite datasets. Our analysis includes three distinct UMAP projections. Each projection provides a unique visual representation of the landscape that highlights the heterogeneity within IgG+ MBCs, IgG+ ActBCs and ASCs.

### Statistics and data visualization

Statistics and data visualization were performed using the programming language R, using RStudio. For the visualization of marker expression, cell frequencies between groups, ggplot2 (V3.3.2), ggpubr (V0.2.5), rstatix (V0.7.0) and ggridges (V0.5.3) packages in R were used. The Wilcoxon signed-rank test was used to compare two or more groups, with unpaired and paired analysis as necessary. The results were adjusted for multiple comparisons using the Holm-Bonferroni correction method as implemented in the rstatix package. The nonparametric Spearman’s rank-order correlation was used to test for correlation. We used the following convention for symbols indicating statistical significance; ns P > 0.05, * P ≤ 0.05, ** P ≤ 0.01, *** P ≤ 0.001, **** P ≤ 0.0001.

## Supplementary Materials

Fig. S1 to S6

Table S1 to S3

## Acknowledgements

We would like to thank all patients and donors who participated in this study. Furthermore, we would like to thank the people of the Sanquin COVID-19 biobank for the collection and processing of samples. In addition, we wish to thank all members of the RECoVERED Study Group, listed below.

RECoVERED Study Group:

Public Health Service of Amsterdam: Ivette Agard, Jane Ayal, Floor Cavdar, Annemarieke Deuring, Annelies van Dijk, Ertan Ersan, Laura del Grande, Joost Hartman, Tjalling Leenstra, Romy Lebbink, Dominique Loomans, Tom du Maine, Ilja de Man, Amy Matser, Lizenka van der Meij, Marleen van Polanen, Maria Oud, Clark Reid, Leeann Storey, Marc van Wijk.

Amsterdam University Medical Centres: Joyce van Assem, Marijne van Beek, Thyra Blankert, Leah Frenkel, Jelle van Haga, Xiaochuan (Alvin) Han, Agnes Harskamp-Holwerda, Mette Hazenberg, Soemeja Hidad, Neeltje Kootstra, Lara Kuijt, Colin Russell, Karlijn van der Straten, Gerben-Rienk Visser.

## Author contributions

Conceptualization: MC, GE, JJGV, AB, MJG, SMH

Methodology: MC, GE, GK, AGAP, TR, JJGV, AB, MJG, SMH

Investigation: MC, GE, GK, LHK, MD, AGAP, JAB, MPo, WO, NJ, RJ

Visualization: MC, GE

Sample acquisition: EW, HDGW, MPr, GJB, MJ, TWK, FE, CES,

Funding acquisition: MC, MJG, SMH

Project administration: MC, GE, MJG, SMH

Supervision: MC, GE, TR, JJGV, MJG, SMH

Writing – original draft: MC, GE, MJG, SMH

Writing – review & editing: All authors

## Data availability

All data is readily available in the main text and supplementary materials. Flow Cytometry Standard (FCS) data generated in this study will be deposited at Zenodo (https://doi.org/10.5281/zenodo.10368326). All reasonable requests for code and materials used in this study should be directed to and will be fulfilled under an MTA by Prof. SM van Ham and Dr. MJ van Gils.

---

**FIGURE S1:**
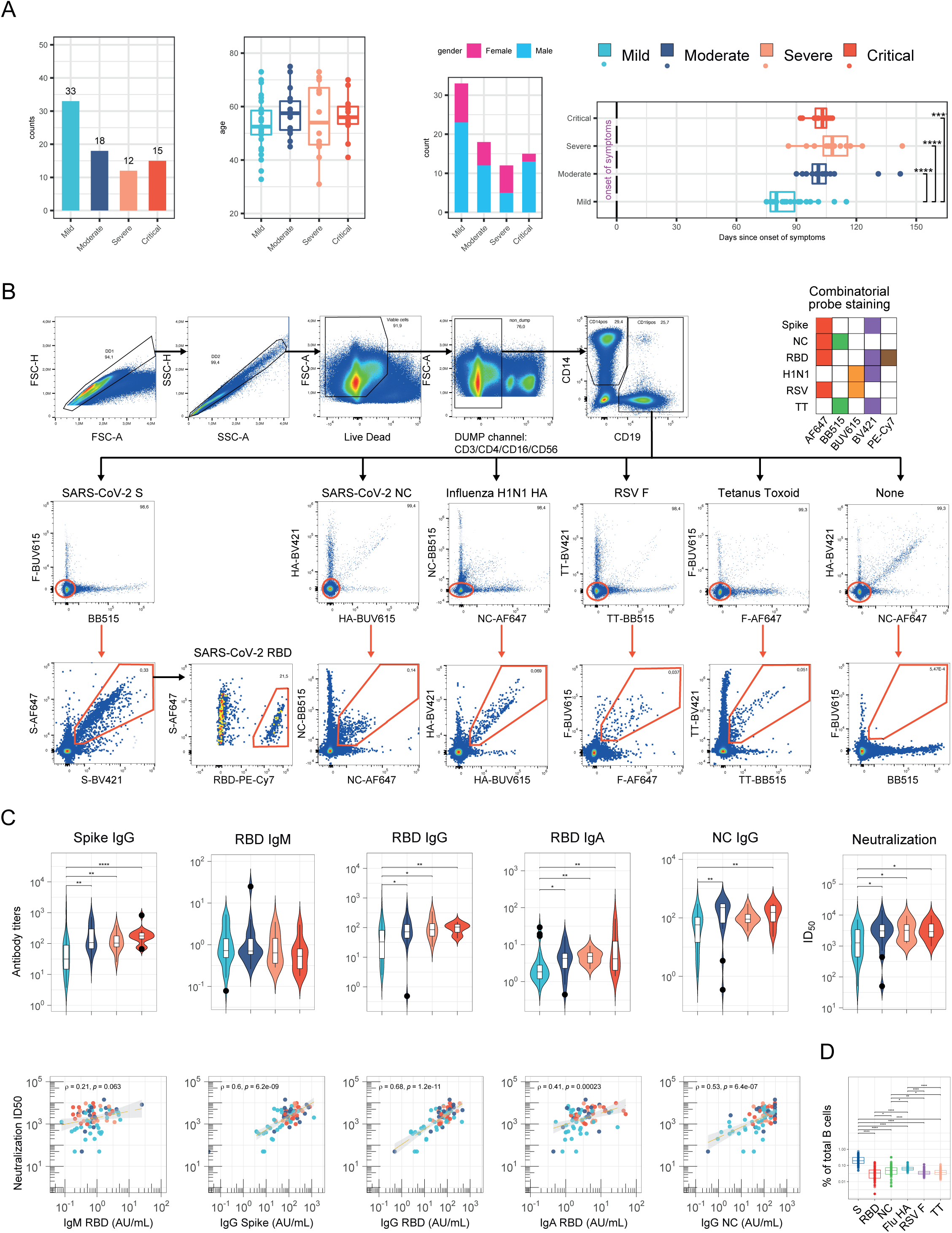
Patient data, sera antibodies and antigen-specific B cells. **(A)** This figure summarizes data from 78 SARS-CoV-2 convalescent patients, categorizing them by disease severity: mild, moderate, severe, and critical categories. (Far Left): Patient counts according to disease severity. (Left): Patient age value distribution within each severity group. (Right): A bar chart displaying gender distribution across severity categories. (Far Right): A time-series revealing symptom onset duration in each severity group. **(B)** Combinatorial probe staining and gating strategy for the detection of multiple B cell specificities in a single PBMC sample (see method section). Top panel, gating strategy to identify live B cells: Doublet, Dead cells, CD3+, CD4+, CD16+ and CD56+ cells were excluded. Middle panel: To remove potential cross-reactive B cells to streptavidin, each probe combination was first gated on cells double negative for the two other probe channels. Bottom panel: Antigen-specific B cells are then detected as double positive for the binding of the same antigen multimerized with two different fluorochromes according to a matrix code. RBD specific B cells are detected out of Spike-specific B cells. **(C)** This figure features violin plots displaying antibody titers (top panel) and neutralization titers (bottom panel) for patients’ sera, categorized by disease severity. Moving from left to right: Spike IgG Titer (Far Left), RBD IgM Titer, RBD IgG Titer, RBD IgA Titer, NC IgG Titer, Neutralization Titer (Far right). **(D)** Frequency of reactive B cells out of total B cells, according to B cell specificity.

**Figure S2:**
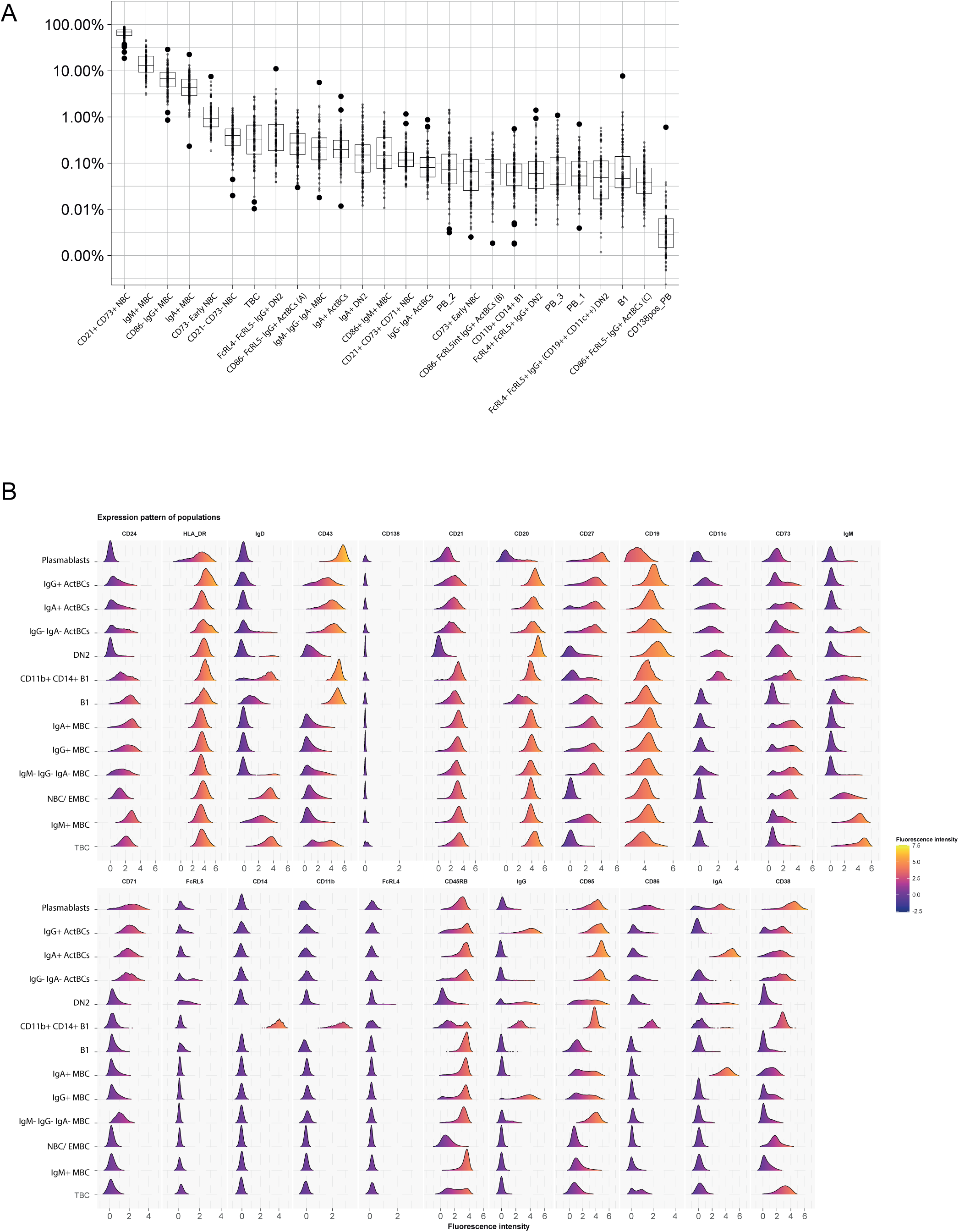
FlowSOM B cell populations. This figure is related to Fig. 1 of the manuscript and provides information about the distribution and phenotype of B cell populations generated by FlowSOM hierarchical clustering based on 23 B cells markers. **(A)** Frequency of 26 B cell populations out of total B cells. **(B)** Comparative analysis of cell surface expression by histogram representation of the 23 B cell markers between the 13 Major B cell populations.

**Figure S3:**
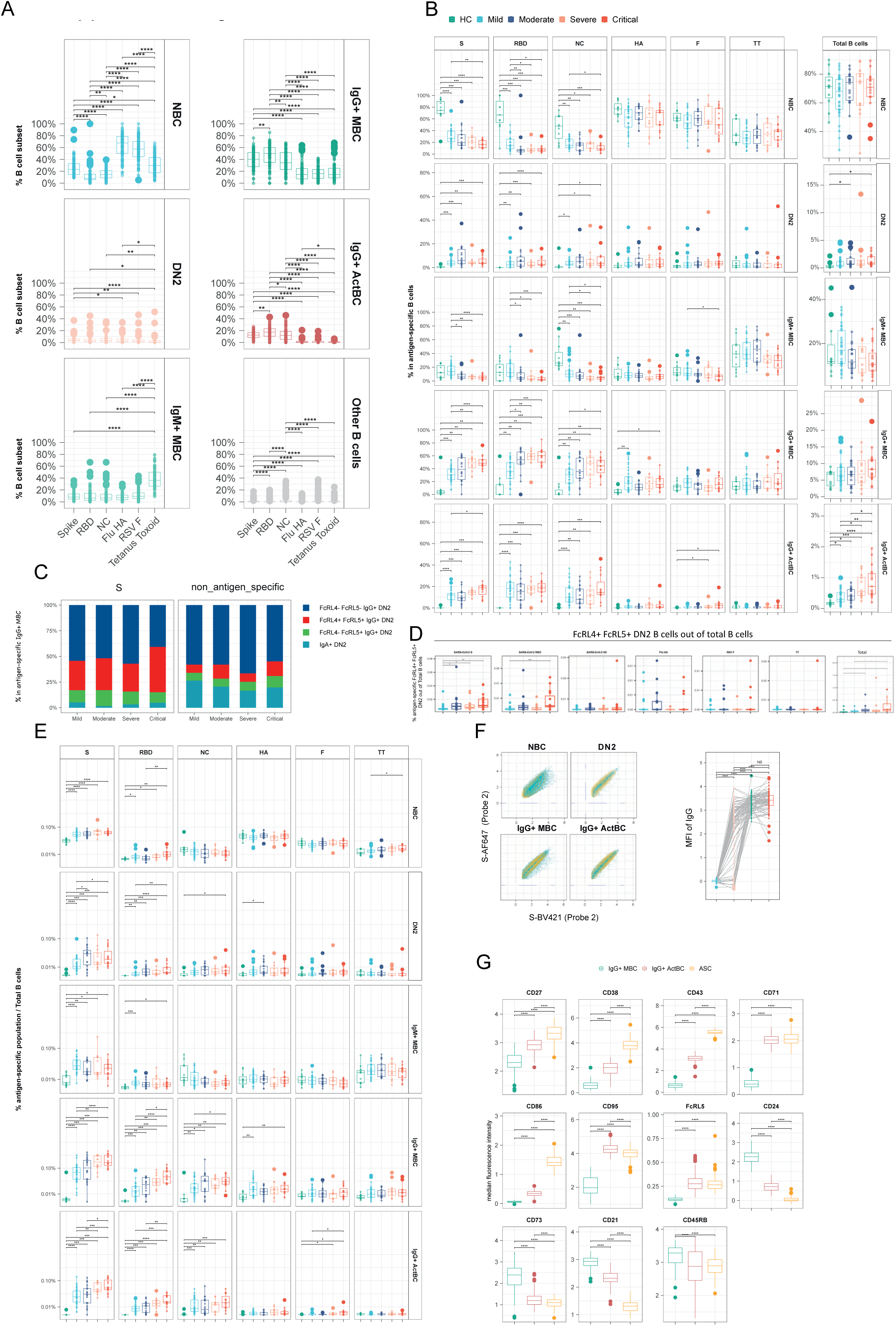
Antigen-specific B cell phenotype. This figure is related to Fig. 1 **(A)** Comparative analysis of B cell subset frequency according to B cell specificity (Spike, RBD, NC, HA, RSV-F, TT, Total B cells) in the five most represented populations Naive, IgM+ Memory B cells, DN B cells, IgG+ memory B cells, IgG+ Activated B cells). **(B)** Frequency of the five most represented populations among each B cell specificity and total B cells, according to disease severity. **(C)** Frequency of DN2 B cell subsets defined by FlowSOM analysis into spike B cell specificity and B cells with no-defined specificity according disease severity. **(D)** Frequency of antigen-specific B cells with FcrL5+ FcrL4+ IgG+ DN2 B cell phenotype in total B cells, according to disease severity. **(E)** Frequency of antigen-specific B cells with Naive, IgM+ Memory B cells, DN B cells, IgG+ memory B cells, or IgG+ Activated B cells phenotype in total B cells, according to B cell specificity and disease severity. **(F)** (Left) FACS plot depicting SARS-CoV-2 Spike binding Median fluorescence intensity with 2 different fluorochrome (Spike-AF647 and Spike-BV421) to reactive B cell that stem from different B cell subsets (Naive, DN, IgG+ MBC and IgG+ ActBC. (Right) Comparative analysis of IgG MFI in these 4 same subsets. **(G)** Comparative analysis of cell surface expression (MFI) by box plots representation of 11 relevant B cells markers between IgG+ memory B cells, IgG+ activated B cells, and Plasmablast major populations.

**Figure S4:**
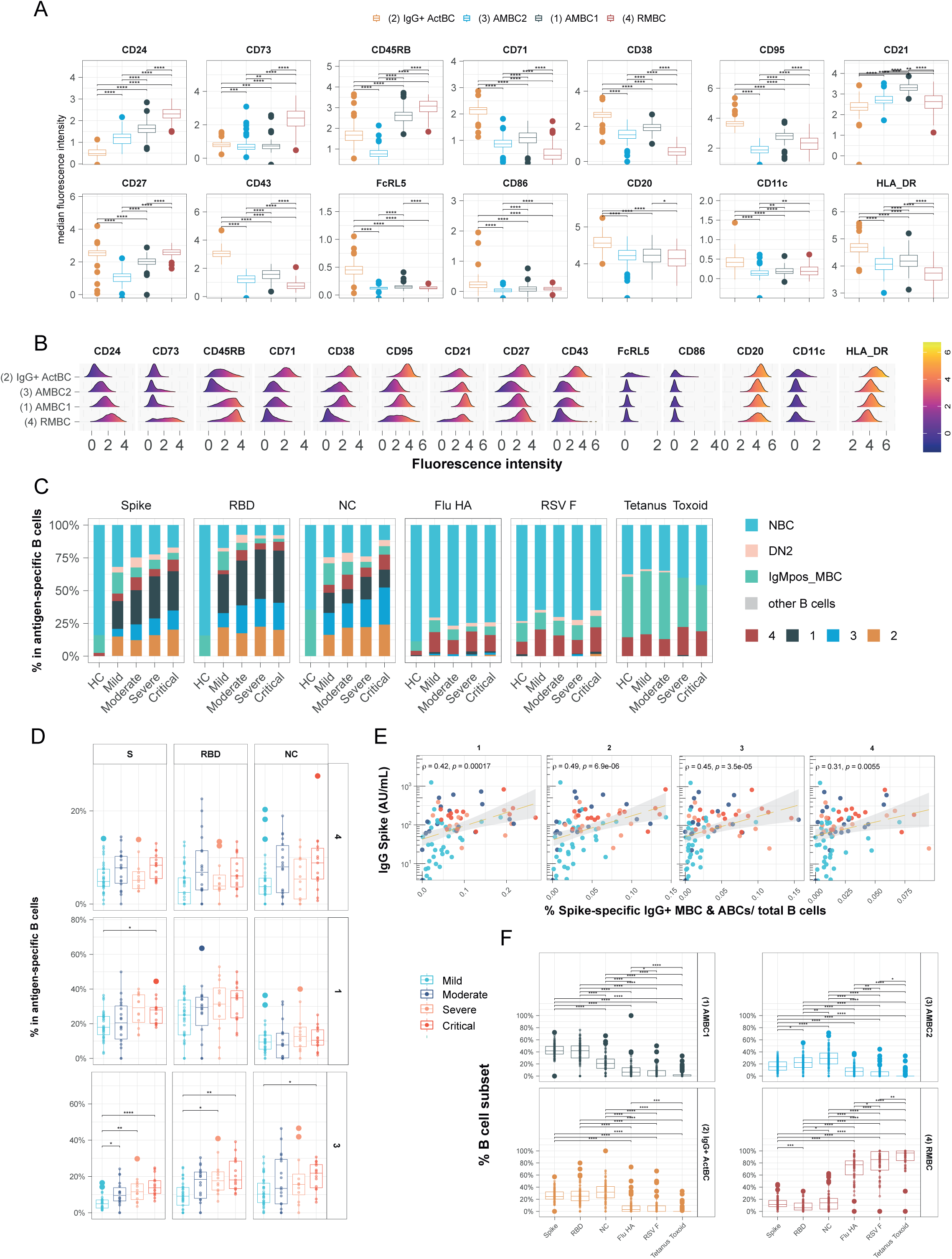
Antigen-specific IgG+ Activated B cells and Memory B cells unsupervised analysis. This figure is related to Fig. 2 Four populations (1: AMBC1 , 2: IgG+ ActBC , 3: AMBC2 , 4: IgG+ RMBC) are comparatively analyzed in this figure and stem from UMAP analysis / and community detection based on Leiden clustering of a composite dataset made of IgG+ Memory B cells and IgG+ Activated B cells that show specificity to any of the 6 antigen studied. **(A)** Comparative analysis of cell surface expression (MFI) by box plots representation of 14 relevant B cells markers between the four populations of interest. **(B)** Comparative analysis of cell surface expression by histogram representation of the 14 relevant B cell markers between the 4 populations of interest. **(C)** Frequency of B cell subsets as identified by FlowSOM-based clustering (Naive, DN2, IgM+ MBC) and Leiden clustering (1: AMBC1 , 2: IgG+ ActBC , 3: AMBC2 , 4: IgG+ MBC) according to B cell specificity and disease severity. **(D)** Frequency of three populations belonging to IgG+ MBC compartment (1: AMBC1 , 3: AMBC2 , 4: IgG+ MBC) among each B cell specificity, according to disease severity. **(E)** Correlation of IgG antibody titer and Neutralization titer with frequencies of the four populations of interest. **(F)** Comparative analysis of B cell subset frequency according to B cell specificity (Spike, RBD, NC, HA, RSV F, TT, Total B cells) in 1: AMBC1 , 2: IgG+ ActBC , 3: AMBC2 , 4: IgG+ MBC.

**Figure S5:**
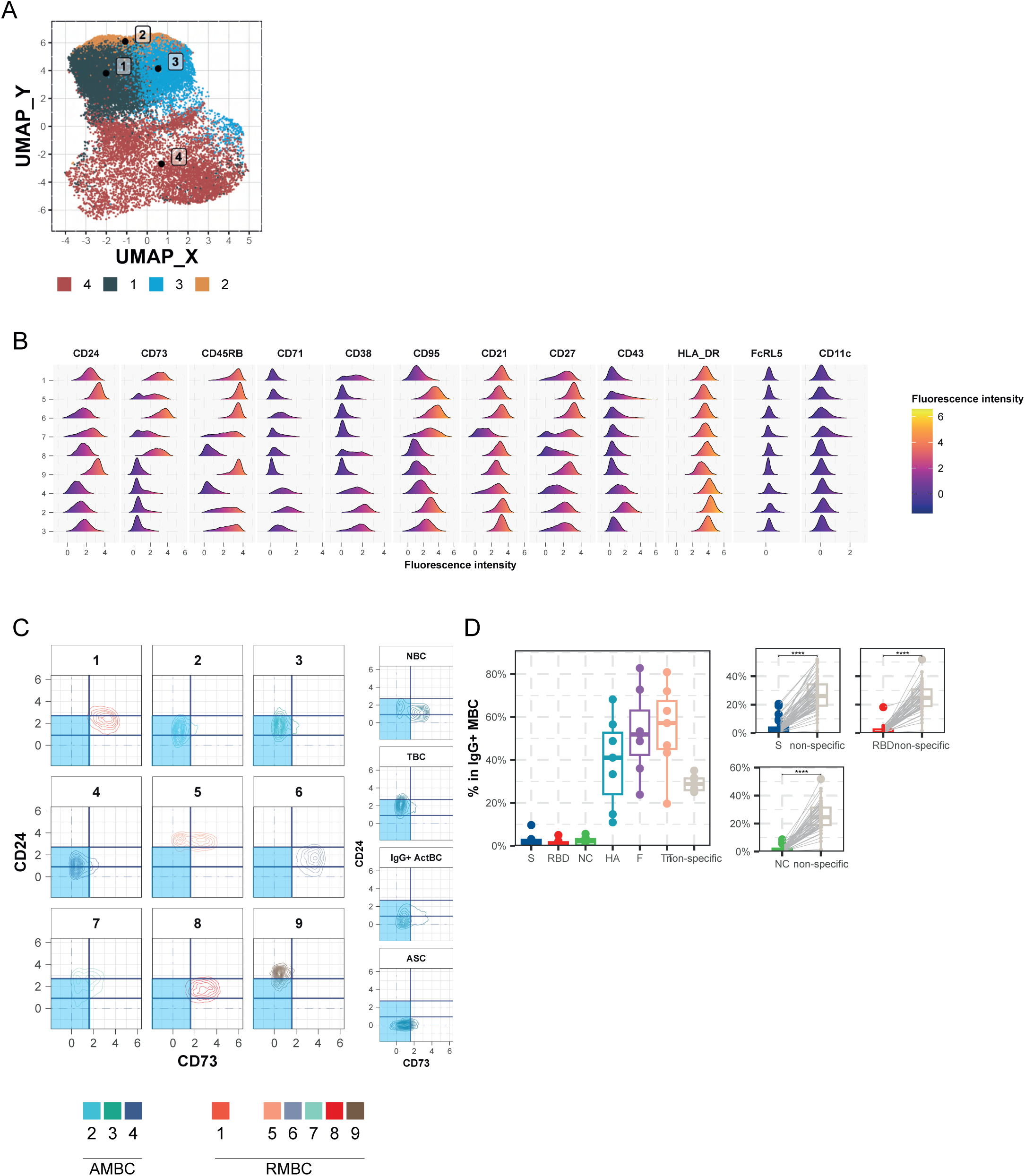
IgG+ memory B cells unsupervised analysis. This figure is related to Fig. 3 Populations analyzed in this figure stem from the IgG+ Memory B cells major population segregated by FlowSOM clustering. From this population a composite dataset has been generated encompassing IgG+ Memory B cells that show specificity to any of the six antigens studied and an additional 1000 non-specific B cells from each individual donor of the IgG+ memory B cells major population. **(A)** Overlay of four IgG+ populations as identified by Leiden clustering in Fig. 2 (1: AMBC1, 2: IgG+ ActBC, 3: AMBC2, 4: IgG+ MBC) displayed on the UMAP data generated out of IgG+ MBC antigen-specific B cells and non-reactive IgG+ MBCs in Fig. 3. **(B)** Comparative analysis of cell surface expression by histogram representation of the 12 relevant B cell markers between the 9 clusters as identified by Leiden clustering in Fig. 3. **(C)** FACS plots depicting CD73 vs CD24 expression of the nine subpopulations that stemmed from IgG+ MBCs from Fig. 3. Naive, Transitional, Activated and Antibody secreting B cells were also depicted as comparative for CD73 and CD24 expression. **(D)** Comparative analysis of cluster 1 frequency, according to specificity, within antigen-specific or non-specific IgG+ MBC. (Left) Paired comparative analysis of B cells with HA, RSV-F, TT, S, RBD and NC or none specificities for samples that encompassed at least 20 cells within the IgG+ MBC compartment for the five tested specificities. (Right) Paired comparative analysis of Spike, RBD, and NC versus non-reactive B cells, for samples that encompass at least 20 cells for a given specificity.

**Figure S6:**
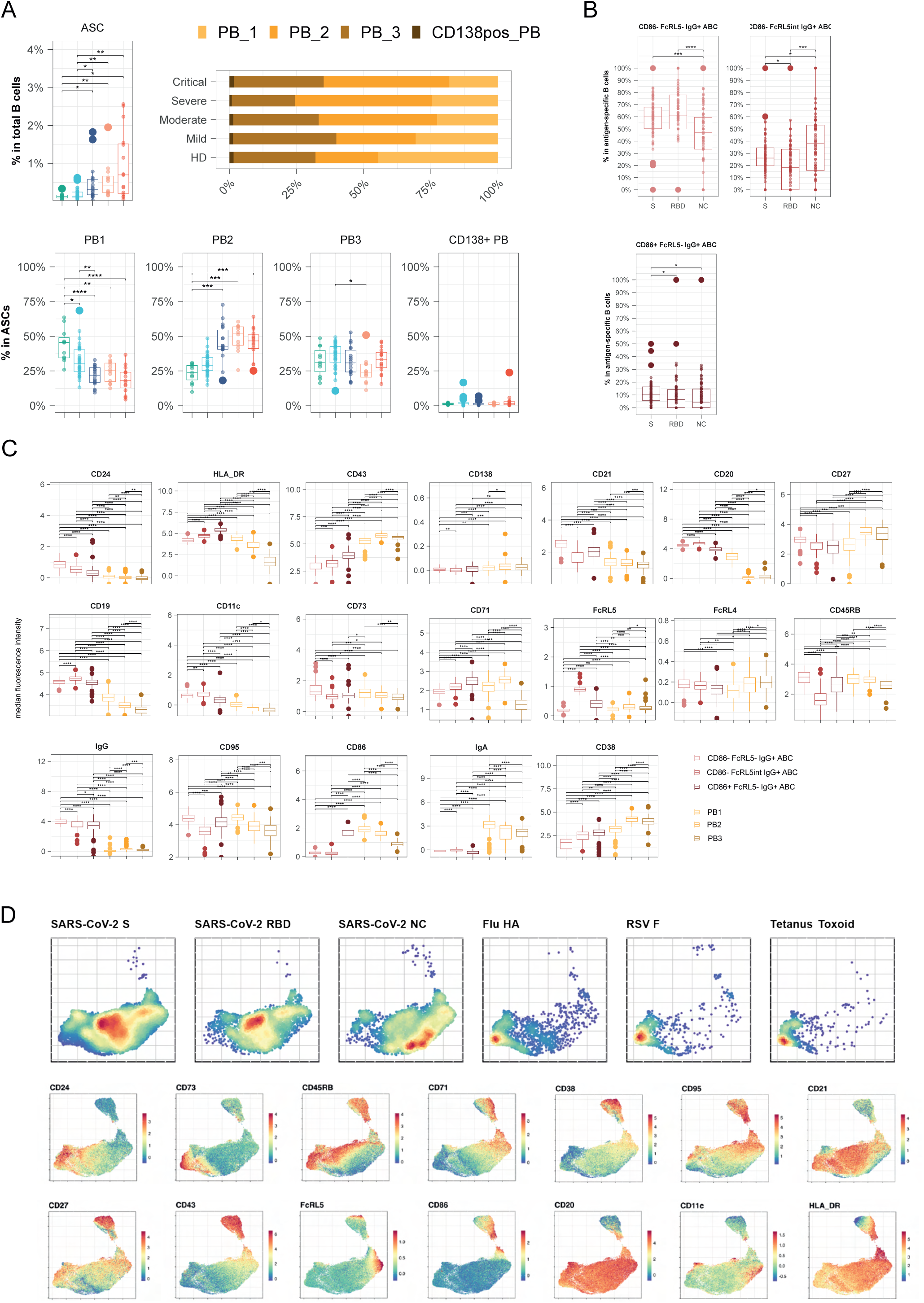
Antibody secreting cells and Activated B cells analysis. This figure is related to Fig. 4 **(A)** (Top left) Frequency of ASC out of total B cells according to disease severity. (Top right & bottom) Frequency of ASC populations (PB_1, PB_2, PB_3, CD138pos_PB) out of total B cells, according to disease severity. **(B)** Frequency of IgG+ activated B cell populations (A: CD86− FcRL5−, B: CD86− FcRL5+, C: CD86+ FcRL5−) out of SARS-CoV-2-specific B cells (S, RBD, NC). **(C)** Comparative analysis of cell surface marker expression (MFI) by box plots representation of 20 relevant B cells markers between plasmablasts populations (PB_1, PB_2, PB_3, CD138pos_PB) and IgG+ activated B cell populations (A: CD86− FcRL5−, B: CD86− FcRL5+, C: CD86+ FcRL5−). **(D)** Feature plots showing scaled normalized counts for 14 relevant B cells markers in a composite data set combining IgG+ MBCs, IgG+ ActBCs and ASCs.

**Table Supplemental 1.**
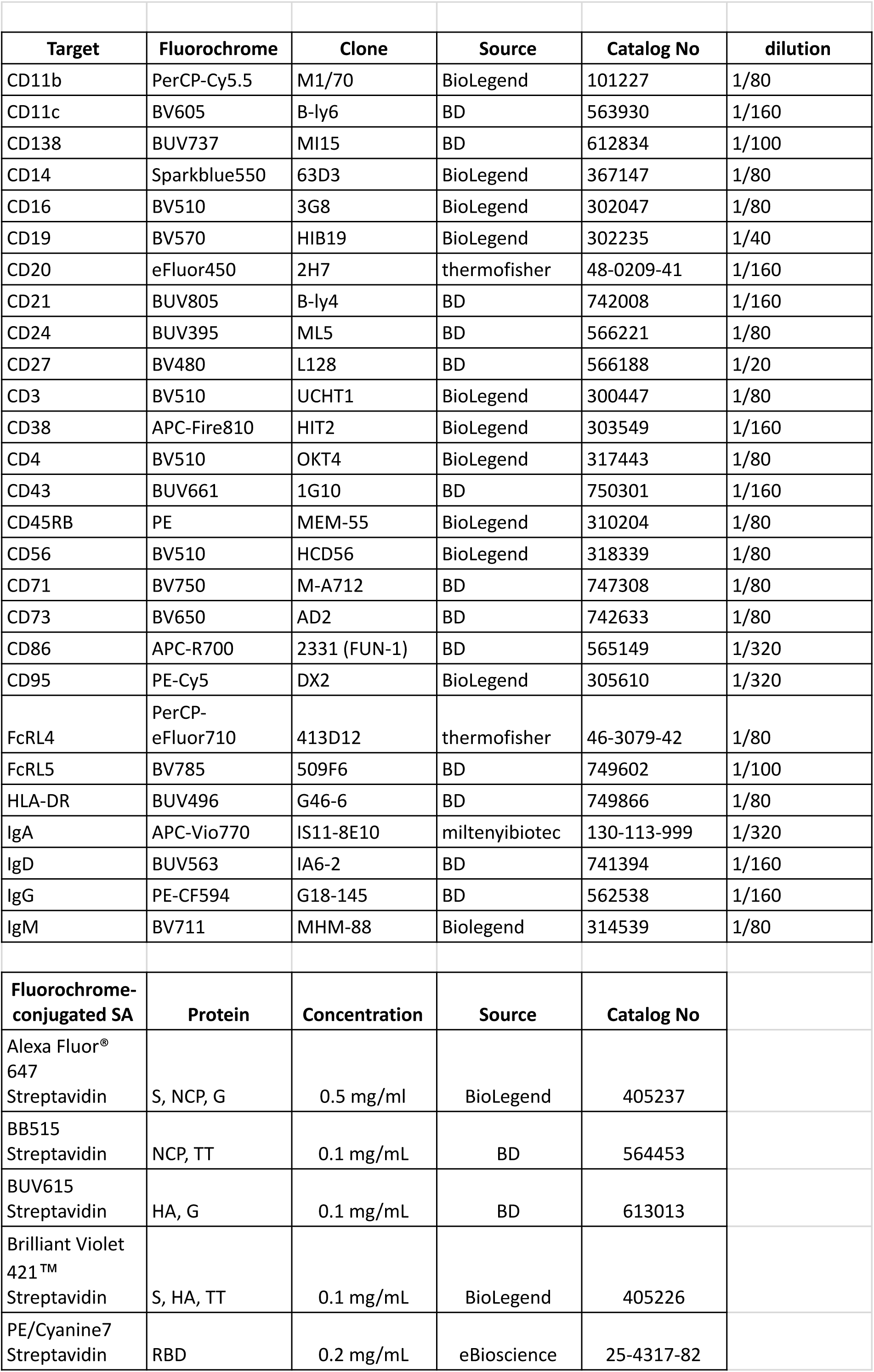

| Target | Fluorochrome | Clone | Source | Catalog No | dilution |
| --- | --- | --- | --- | --- | --- |
| CD11b | PerCP-Cy5.5 | M1/70 | BioLegend | 101227 | 1/80 |
| CD11c | BV605 | B-ly6 | BD | 563930 | 1/160 |
| CD138 | BUV737 | MI15 | BD | 612834 | 1/100 |
| CD14 | Sparkblue550 | 63D3 | BioLegend | 367147 | 1/80 |
| CD16 | BV510 | 3G8 | BioLegend | 302047 | 1/80 |
| CD19 | BV570 | HIB19 | BioLegend | 302235 | 1/40 |
| CD20 | eFluor450 | 2H7 | thermofisher | 48-0209-41 | 1/160 |
| CD21 | BUV805 | B-ly4 | BD | 742008 | 1/160 |
| CD24 | BUV395 | ML5 | BD | 566221 | 1/80 |
| CD27 | BV480 | L128 | BD | 566188 | 1/20 |
| CD3 | BV510 | UCHT1 | BioLegend | 300447 | 1/80 |
| CD38 | APC-Fire810 | HIT2 | BioLegend | 303549 | 1/160 |
| CD4 | BV510 | OKT4 | BioLegend | 317443 | 1/80 |
| CD43 | BUV661 | 1G10 | BD | 750301 | 1/160 |
| CD45RB | PE | MEM-55 | BioLegend | 310204 | 1/80 |
| CD56 | BV510 | HCD56 | BioLegend | 318339 | 1/80 |
| CD71 | BV750 | M-A712 | BD | 747308 | 1/80 |
| CD73 | BV650 | AD2 | BD | 742633 | 1/80 |
| CD86 | APC-R700 | 2331 (FUN-1) | BD | 565149 | 1/320 |
| CD95 | PE-Cy5 | DX2 | BioLegend | 305610 | 1/320 |
| FcRL4 | PerCP-eFluor710 | 413D12 | thermofisher | 46-3079-42 | 1/80 |
| FcRL5 | BV785 | 509F6 | BD | 749602 | 1/100 |
| HLA-DR | BUV496 | G46-6 | BD | 749866 | 1/80 |
| IgA | APC-Vio770 | IS11-8E10 | miltenyibiotec | 130-113-999 | 1/320 |
| IgD | BUV563 | IA6-2 | BD | 741394 | 1/160 |
| IgG | PE-CF594 | G18-145 | BD | 562538 | 1/160 |
| IgM | BV711 | MHM-88 | Biolegend | 314539 | 1/80 |
| Fluorochrome-conjugated SA | Protein | Concentration | Source | Catalog No |  |
| Alexa Fluor® 647 Streptavidin | S, NCP, G | 0.5 mg/ml | BioLegend | 405237 |  |
| BB515 Streptavidin | NCP, TT | 0.1 mg/mL | BD | 564453 |  |
| BUV615 Streptavidin | HA, G | 0.1 mg/mL | BD | 613013 |  |
| Brilliant Violet 421™ Streptavidin | S, HA, TT | 0.1 mg/mL | BioLegend | 405226 |  |
| PE/Cyanine7 Streptavidin | RBD | 0.2 mg/mL | eBioscience | 25-4317-82 |  |

**Table Supplemental 2.**
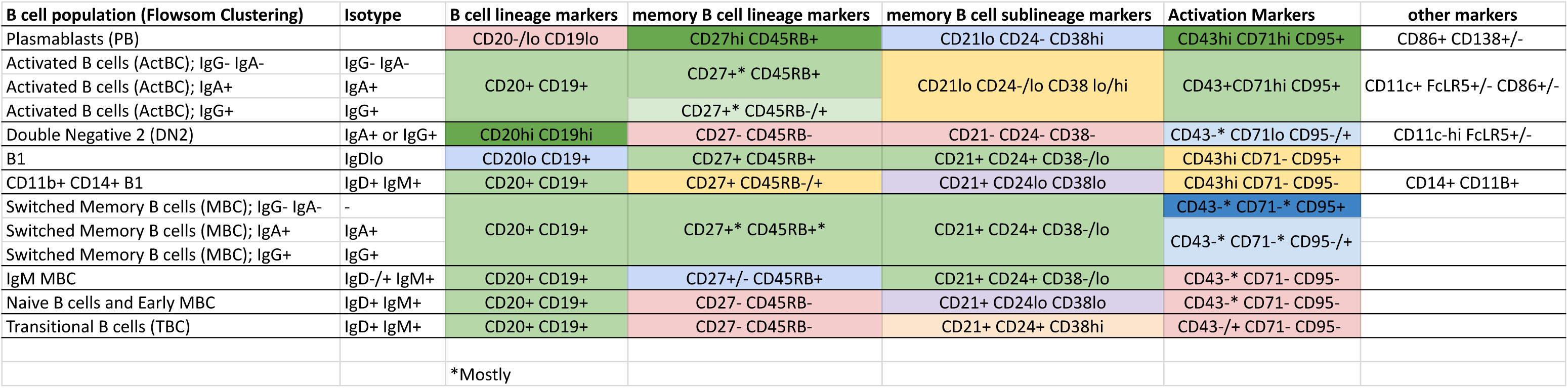

**Supplementary Table 3A.** Related to Figure 2. Most variable markers in antigen-specific IgG+ MBCs and IgG+ ActBC

| marker | Variance | Min. | 1st Qu. | Median | 3rd Qu. | Max. |
| --- | --- | --- | --- | --- | --- | --- |
| CD43 | 1.357 | -0.810 | 0.759 | 1.589 | 2.532 | 6.364 |
| CD45RB | 1.267 | -0.805 | 1.047 | 2.033 | 2.957 | 4.654 |
| CD95 | 1.140 | -0.824 | 1.877 | 2.662 | 3.406 | 5.941 |
| CD27 | 0.884 | -1.024 | 1.409 | 2.130 | 2.736 | 4.935 |
| CD73 | 0.879 | -0.686 | 0.467 | 0.804 | 1.470 | 5.153 |
| CD21 | 0.867 | -0.962 | 2.221 | 2.896 | 3.380 | 5.674 |
| CD38 | 0.861 | -0.924 | 1.026 | 1.782 | 2.370 | 4.567 |
| CD24 | 0.773 | -0.909 | 0.713 | 1.341 | 1.958 | 4.706 |
| CD71 | 0.563 | -0.499 | 0.657 | 1.174 | 1.762 | 4.140 |
| CD20 | 0.360 | 0.265 | 3.824 | 4.223 | 4.597 | 6.183 |
| CD11c | 0.155 | -0.865 | -0.012 | 0.202 | 0.450 | 3.429 |
| CD86 | 0.121 | -0.664 | -0.045 | 0.090 | 0.253 | 4.138 |
| FcRL5 | 0.070 | -0.964 | 0.071 | 0.156 | 0.284 | 2.009 |

**Supplementary Table 3B.** Related to Figure 3. Most variable markers in antigen-specific and non-specific IgG+ MBCs

| marker | Variance | Min. | 1st Qu. | Median | 3rd Qu. | Max. |
| --- | --- | --- | --- | --- | --- | --- |
| CD73 | 1.623 | -0.686 | 0.561 | 1.429 | 2.804 | 5.584 |
| CD45RB | 1.478 | -0.910 | 1.465 | 2.805 | 3.455 | 4.654 |
| CD95 | 1.430 | -1.099 | 1.304 | 2.245 | 3.142 | 5.929 |
| CD27 | 1.106 | -1.067 | 1.280 | 2.153 | 2.834 | 4.881 |
| CD43 | 0.901 | -0.810 | 0.331 | 0.874 | 1.651 | 6.253 |
| CD24 | 0.853 | -0.909 | 1.255 | 1.909 | 2.608 | 4.902 |
| CD38 | 0.818 | -0.939 | 0.244 | 1.067 | 1.845 | 4.143 |
| CD21 | 0.802 | -0.962 | 2.370 | 2.957 | 3.394 | 5.674 |
| CD20 | 0.353 | 0.265 | 3.626 | 4.027 | 4.405 | 6.146 |
| CD71 | 0.327 | -0.564 | 0.250 | 0.600 | 1.100 | 3.644 |
| CD11c | 0.130 | -0.896 | -0.065 | 0.139 | 0.364 | 2.837 |
| CD86 | 0.036 | -0.664 | -0.036 | 0.062 | 0.178 | 1.592 |
| FcRL5 | 0.016 | -0.964 | 0.057 | 0.120 | 0.194 | 1.020 |

**Supplementary Table 3C.**
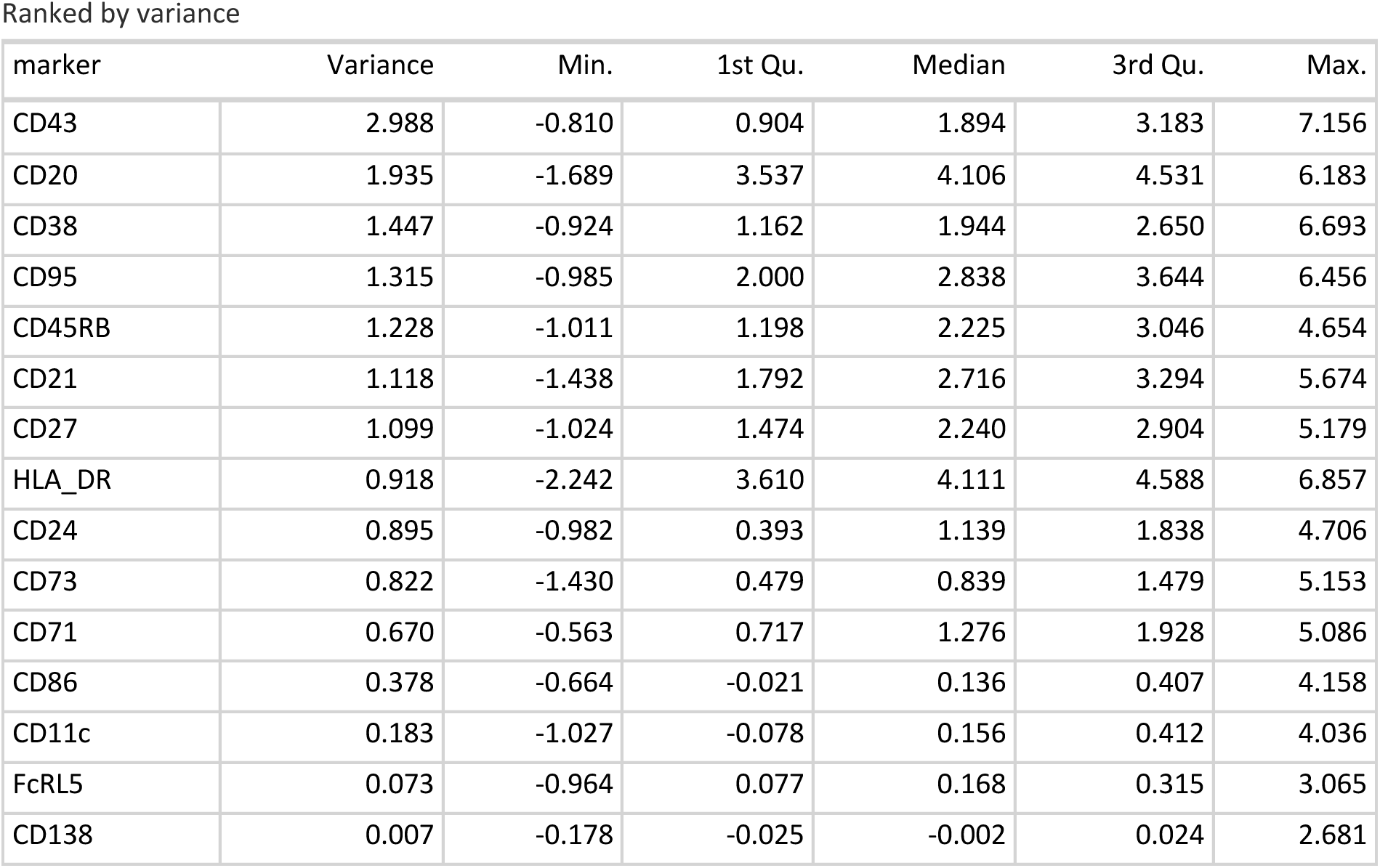
Related to Figure 4. Most variable markers in antigen-specific IgG+ MBCs and IgG+ ActBC and ASC

